# Nanoscale analysis of functionally diverse glutamatergic synapses in the neocortex reveals input and layer-specific organization

**DOI:** 10.1101/2024.05.01.592008

**Authors:** Grace Jones, Yeasmin Akter, Victoria Shifflett, Martin Hruska

**Author notes:** **Corresponding author:** Martin Hruska. Authors contributed equally to this work (Grace Jones, Yeasmin Akter). **Author contributions:** G.J. conducted experiments, analyzed data, designed experiments and wrote the paper; Y.A. conducted experiments, analyzed data, designed experiments and wrote the paper; V.S. analyzed data; M.H. conceptualized the study, designed experiments, conducted experiments, analyzed data and wrote the paper.

## Abstract

Discovery of synaptic nanostructures suggests a molecular logic for the flexibility of synaptic function. We still have little understanding of how functionally diverse synapses in the brain organize their nanoarchitecture due to challenges associated with super-resolution imaging in complex brain tissue. Here, we characterized single-domain camelid nanobodies for the 3D quantitative multiplex imaging of synaptic nano-organization in 6 µm brain cryosections using STED nanoscopy. We focused on thalamocortical (TC) and corticocortical (CC) synapses along the apical-basal axis of layer 5 pyramidal neurons as models of functionally diverse glutamatergic synapses in the brain. Spines receiving TC input were larger than CC spines in all layers examined. However, TC synapses on apical and basal dendrites conformed to different organizational principles. TC afferents on apical dendrites frequently contacted spines with multiple aligned PSD-95/Bassoon nanomodules, which are larger. TC spines on basal dendrites contained mostly one aligned PSD-95/Bassoon nanocluster. However, PSD-95 nanoclusters were larger and scaled with spine volume. The nano-organization of CC synapses did not change across cortical layers. These results highlight striking nanoscale diversity of functionally distinct glutamatergic synapses, relying on afferent input and sub-cellular localization of individual synaptic connections.

## Introduction

Synapses in the central nervous system are small, diffraction-limited cell-cell junctions designed to rapidly transfer and process information. One characteristic feature of central synapses is their remarkable structural and functional variability [1, 2]. A single pyramidal neuron receives thousands of synapses that differ in strength, neurotransmitter release probability and signaling properties that form the basis for cortical computation and information storage in the brain [3–11]. At each synapse, diverse proteins with distinct lifetimes organize at the nanoscale in an ordered manner to build active zones and post-synaptic densities (PSDs) that endow synapses with exquisite regulation of synaptic function [12–15]. Despite the importance of nanoscale ensembles for brain development, learning and memory, and aging, how functionally diverse synapses organize their pre- and post-synaptic nano-architecture and how the source of afferent input might shape synaptic nano-organization is not understood [16].

Super-resolution imaging of synapses *in vitro* demonstrated that structural and functional ensembles exhibit conserved nano-organization where precisely aligned nanoclusters of pre- and post-synaptic proteins occupy discrete positions in PSDs and active zones [17–25]. In dendritic spines, the number of aligned trans-synaptic units, not their size, scales with the increasing size of dendritic spines [18, 19]. This modular nano-architecture was suggested to provide flexibility needed for events such as synaptic plasticity [26–29]. Both structural and functional constituents of the synapse, including PSD-95, AMPARs, NMDARs and presynaptic Bassoon, VGluT1, and Synaptotagmin-1 conformed to this scalability, suggesting the robustness of modular nano-organization [18, 19]. Understanding how these nanoscale principles translate to the organization of functionally diverse synapses in the brain will require 3D reconstruction of molecularly identified connections in their native environment.

Pyramidal neurons in the somatosensory cortex receive functionally diverse glutamatergic innervation from corticocortical (CC) and thalamocortical (TC) synapses. TC synapses account for the minority (5-20%) of all synapses in the cortex yet are highly efficient in activating cortical neurons [30–35]. How sparse TC innervation can efficiently drive cortical excitation has been debated. *Ex vivo* studies suggested that TC synapses onto spiny neurons in layer 4 (L4) are multi-quantal and several-fold stronger than unitary CC synaptic connections onto the same neurons [36–38]. Others demonstrated that TC and CC synapses in L4 are indistinguishable electrophysiologically; instead, clustering of TC synapses in specific dendritic domains was suggested to synchronize feedforward activity [39–41]. More recently, work using expansion microscopy in the mouse visual cortex showed that spines of L2/3 pyramidal neurons receiving TC input are smaller and weaker than neighboring CC synapses [42]. These data suggest that mechanisms regulating TC synaptic function are complex and vary depending on the cell type and cortical location. Understanding the principles of nano-organization of TC and CC synapses across cortical layers will shed light on processing of cortical information in the brain.

Super-resolution imaging in complex brain tissue is challenging. The small size of synapses combined with the protein-rich environment of active zones and PSDs make it difficult for antibodies to efficiently label molecules in the aldehyde fixed tissue [14, 43]. Furthermore, resolution decreases with depth, limiting super-resolution only to the first few microns from the tissue surface [44]. Single-domain camelid nanobodies that are much smaller than conventional antibodies are promising tools for facilitating super-resolution imaging in brain tissue [45–48]. The efficacy of nanobodies for quantitative multiplex super-resolution imaging of endogenous synaptic proteins *in situ* needs to be determined.

Using Stimulated Emission Depletion (STED) nanoscopy, we show that labeling with nanobodies reduces errors in synapse identification. Moreover, nanobodies outperform antibodies in obtaining the sub-diffraction resolution needed to quantify synaptic nano-organization *in situ*. Combining nanobody labeling with five-channel confocal and 3D tau-STED imaging of spines on L5 pyramidal neurons, we demonstrate that spines receiving TC synapses are larger than CC spines on both apical and basal dendrites. However, the nano-organization of TC synapses changes across the apical/basal axis. TC spines in apical tufts contain multiple trans-synaptic nanoclusters and conform to modular principles of nano-organization. In contrast, TC spines on basal dendrites contain predominantly one trans-synaptic PSD-95/Bassoon nanocluster. In basal spines, nanocluster sizes scale with spine volume, suggesting that they are organized in a non-modular manner. Our findings highlight the importance of afferent input and subcellular localization in shaping the nano-architecture of functionally diverse glutamatergic synapses.

## Results

### Conventional antibodies are inefficient for the visualization of synapses *in situ*

A key requirement for 3D nano-scale reconstructions of individual synapses in complex brain tissue is the ability to accurately localize pre- and post-synaptic proteins to the same spine. However, immunostaining for many synaptic molecules, including PSD-95, has been shown to produce relatively high false negative rates [49]. Therefore, colocalization of fluorescently labeled spines with only a pre-synaptic marker has been used for *in vivo* synaptic mapping and reconstructions [35, 40]. These approaches exhibit variable accuracy of synapse detection.

To identify putative excitatory synapses in the brain, we labeled spines of L5 pyramidal cells with antibodies for PSD-95 (PSDs), VGluT1 (synaptic vesicles) and Bassoon (active zones) in 50 µm brain sections (Fig. 1). We collected brain sections from the somatosensory cortex (S1) of Thy-1-YFP-H mice in which L5 pyramidal neurons express high levels of GFP spectral variant yellow fluorescent protein (YFP) [50]. Brain sections were either directly used for immunolabeling with the above antibodies or subjected to antigen retrieval (AR), a CUBIC tissue clearing approach, or CUBIC + AR before the antibody labeling (Fig. 1a) [19, 51]. We measured synaptic puncta density along the YFP-labeled primary and secondary branches in apical tufts of L5 pyramidal neurons. PSD-95 puncta density was not significantly different between various treatments (Fig. 1b). Using average PSD-95 puncta density measurements from each condition, we randomized PSD-95 clusters using the Monte Carlo (MC) macro in ImageJ [19]. Surprisingly, the density of PSD-95 was not significantly higher than MC simulations regardless of the tissue treatment (Fig. 1b).

**Figure 1.**
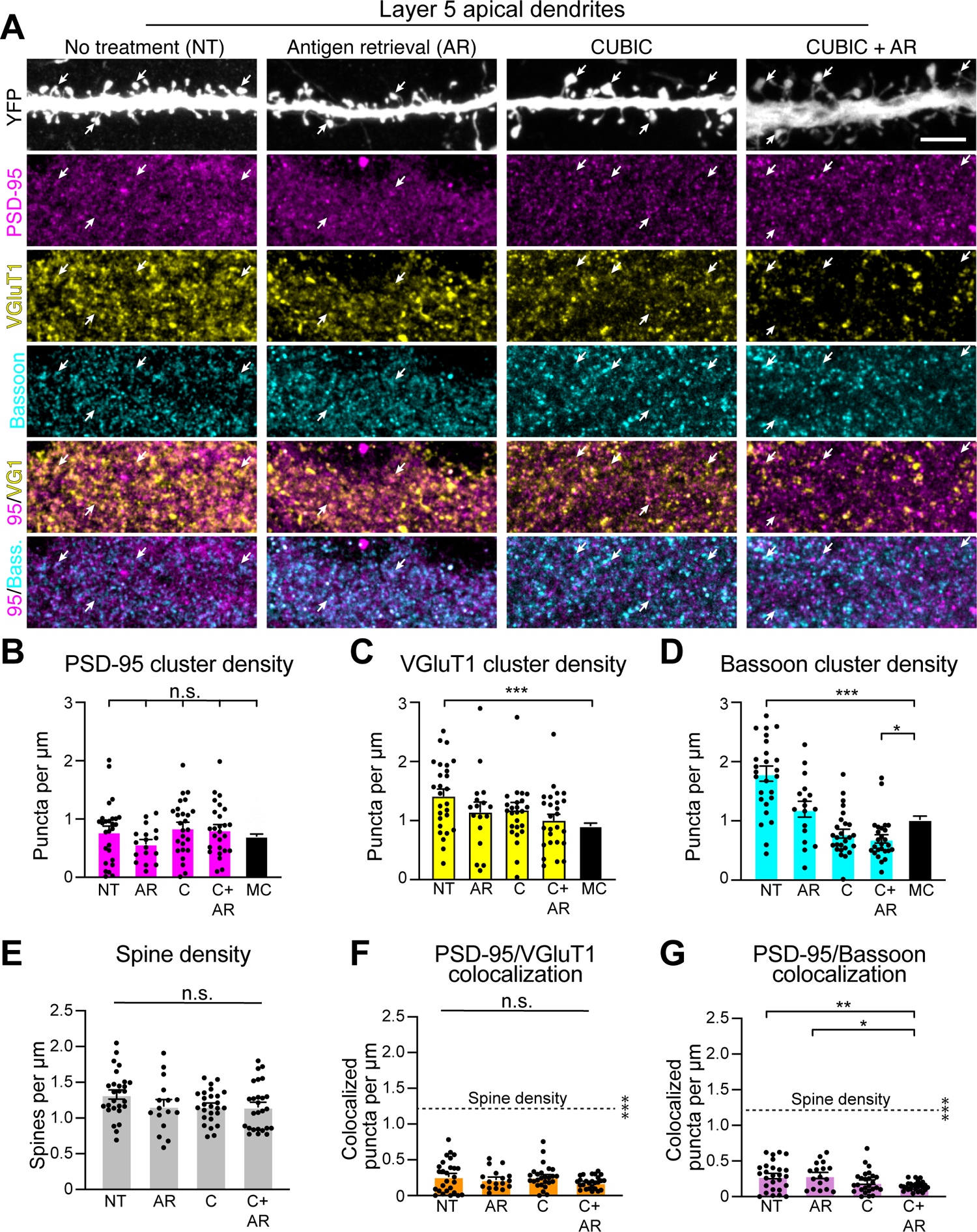
Conventional antibodies inefficiently label synapses in brain sections. **(A)** Representative confocal images of L5 apical dendrites in 50 µm cortical sections collected from Thy1-YFP-H mice. Brain sections were either directly used for immunolabeling with antibodies against PSD-95 (magenta), VGluT1 (yellow) and Bassoon (cyan) to visualize pre- and post-synaptic specializations on YFP-labeled dendrites (gray, enhanced by GFP antibody) or subjected to antigen retrieval and CUBIC clearing before immunolabeling. Arrows indicate PSD-95 labeling in dendritic spines. Scale bar, 5 µm. **(B-D)** Quantification of average PSD-95 (**B**), VGluT1 (**C**), and Bassoon (**D**) densities along YFP-labeled apical dendrites. Average cluster densities in untreated (NT, n = 27 neurons), Antigen Retrieved (AR, n = 17 neurons), CUBIC (C, n = 26 neurons) and CUBIC+Antigen Retrieved (C+AR, n = 26 neurons) conditions were compared to densities of Monte Carlo (MC) randomized clusters for PSD-95 (p = 0.1439), VGluT1 (***p<0.0001) and Bassoon (***p<0.0001, *p = 0.015, ANOVA, Dunnett’s post-hoc). Randomizations were performed using experimentally derived cluster numbers from all antibodies in all treatment conditions. **(E)** Quantification of dendritic spine densities in apical dendrites of L5 neurons in the primary somatosensory cortex (p = 0.1429, ANOVA). **(F, G)** Quantification of colocalized PSD-95 and VGluT1 (p = 0.2481) or PSD-95 and Bassoon (**p = 0.0072, *p = 0.0131, ANOVA, Tukey’s post-hoc) cluster densities in apical dendrites of L5 neurons. Average spine density (dotted lines in **F** and **G**) comparisons to colocalized clusters in **F** (***p<0.0001) and **G** (***p<0.0001, ANOVA, Dunnett’s post-hoc). Bar graphs represent the mean ± SEM. Data were collected from a minimum of ten different neurons (dots on graph) from brain sections acquired from at least three different male or female Thy1-YFP-H mice.

Next, we quantified the labeling of pre-synaptic specializations using antibodies against VGlutT1 and Bassoon (Fig. 1c, d). VGluT1 puncta density was higher on average than PSD-95 puncta density. However, only the density of VGluT1 puncta in non-treated brain sections was significantly higher than MC simulations (Fig. 1c). We saw similar results by staining for the active zone marker Bassoon (Fig. 1d). These data indicate that while immunostaining for pre-synaptic proteins is more efficient than labeling of PSD-95, antigen retrieval and CUBIC treatments of brain tissue do not necessarily improve the efficacy of synaptic protein labeling in 50 µm brain sections.

To determine the accuracy of excitatory synapse identification using antibodies, we measured the density of colocalized PSD-95 with either VGluT1 or Bassoon puncta. We compared it to the density of dendritic spines in the apical tufts of L5 neurons.

Dendritic spine density was not different between different tissue treatments (Fig. 1e). However, densities of colocalized PSD-95/VGluT1 and PSD-95/Bassoon clusters were significantly smaller than spine densities (Fig. 1f, g). Assuming that most spines contain a functional synapse [3], colocalized pre- and post-synaptic antibody markers identified only a small fraction of excitatory synapses (∼19% for PSD-95/VGluT1 and ∼ 22% for PSD-95/Bassoon). These data demonstrate that simultaneous labeling for pre- and post-synaptic markers in brain tissue underestimates the number of excitatory synapses by a large margin.

The inefficient synaptic labeling could be due to the inability of antibodies to penetrate the thickness of 50 µm brain sections. Therefore, we collected 6 µm cryosections from S1 of Thy-1-YFP-H mice. As above, we immunostained for PSD-95, VGluT1 and Bassoon either without any treatment or following AR and CUBIC (Fig. 2a). Untreated and CUBIC-only treated sections were still not significantly different from MC simulations (Fig. 2b). However, antigen retrieval alone or in combination with CUBIC significantly increased PSD-95 puncta density over noise (Fig. 2b).

**Figure 2.**
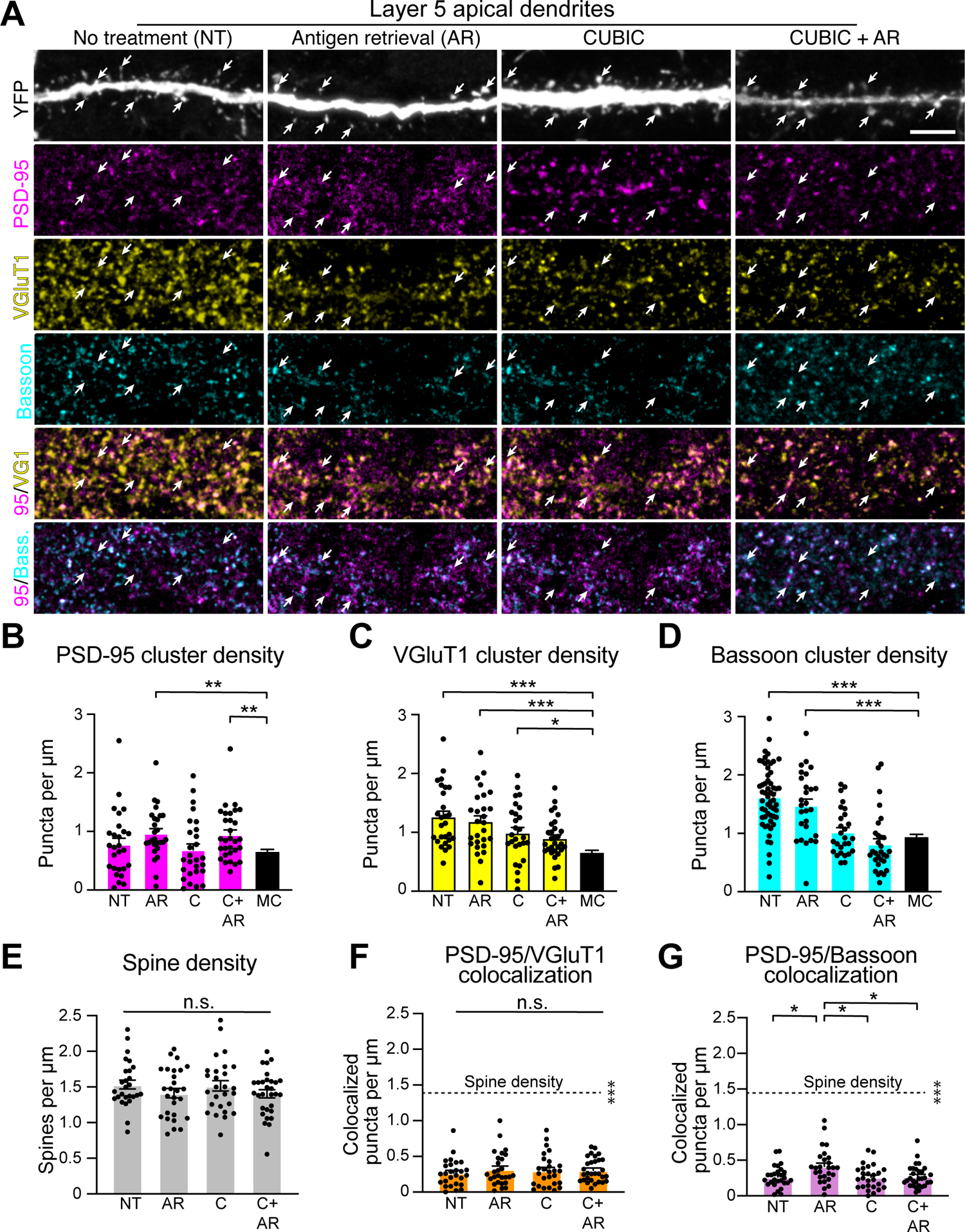
Labeling of synaptic clusters by conventional antibodies is marginally improved in thin cryosections. **(A)** Representative confocal images of L5 apical dendrites in 6 µm cortical cryosections collected from Thy1-YFP-H mice. Cryosections were either directly used for immunolabeling with antibodies against PSD-95 (magenta), VGluT1 (yellow) and Bassoon (cyan) to visualize pre- and post-synaptic specializations on YFP-labeled dendrites (gray, enhanced by GFP antibody) or subjected to antigen retrieval and CUBIC clearing before immunolabeling. Arrows indicate PSD-95 labeling in dendritic spines. Scale bar, 5 µm. **(B-D)** Quantification of PSD-95 (**B**), VGluT1 (**C**) and Bassoon (**D**) densities along YFP-labeled apical dendrites. Average cluster densities in No Treatment (NT, n = 27 neurons), Antigen Retrieved (AR, n = 27 neurons), CUBIC (C, n = 27 neurons) and CUBIC+Antigen Retrieved (C+AR, n = 30 neurons) conditions were compared to densities of Monte Carlo (MC) randomized clusters for PSD-95 (**p<0.005), VGluT1 (***p<0.0001, *p = 0.0229) and Bassoon (***p<0.0001, ANOVA, Dunnett’s post-hoc). Randomizations were performed using experimentally derived cluster numbers from all antibodies in all treatment conditions. **(E)** Quantification of dendritic spine densities in apical dendrites of L5 neurons in the primary somatosensory cortex (p = 0.5556, ANOVA). **(F, G)** Quantification of colocalized PSD-95 and VGluT1 (p = 0.7983) or PSD-95 and Bassoon (*p<0.05, ANOVA, Tukey’s post-hoc) cluster densities along the apical shafts of layer 5 neurons. Average spine density (dotted lines from **e**) comparisons to colocalized clusters in **F** (***p<0.0001) and **G** (***p<0.0001, ANOVA, Dunnett’s post-hoc). Bar graphs represent the mean ± SEM. For each condition, data were collected from a minimum of ten different neurons (dots on graph) from brain sections acquired from at least three different male or female Thy1-YFP-H mice.

Labeling of pre-synaptic clusters was also improved. The density of VGluT1 clusters was significantly higher than Monte Carlo simulations in untreated, AR and CUBIC-treated sections (Fig. 2c). Similarly, Bassoon densities were significantly higher than a random Bassoon cluster distribution in untreated and AR sections but not in CUBIC-cleared sections either with or without AR (Fig. 2d). Spine densities in L5 neuron apical tufts were similar between different treatments and resembled spine densities in 50 µm sections (Fig. 2e). However, colocalized PSD-95/VGluT1 or PSD-95/Bassoon clusters still identified only 17% and 25% of putative excitatory synapses respectively, than what would be expected if all spines contained a functional contact (Fig. 2f, g). Altogether, these data indicate that identifying endogenous proteins with conventional antibodies underestimates the number of excitatory synapses *in situ*.

### Single-domain nanobodies improve the efficiency of synaptic labeling in brain sections

We next sought to characterize reagents that might be better suited for visualizing and quantifying excitatory synapses in their native environment. Single-domain nanobodies are 15kDa molecules derived from camelid heavy chain only antibodies that are ∼ 4 nm in length and 2.5 nm in width [46]. Due to their small size, they have the potential to better penetrate the highly proteinaceous environment of PSDs in brain sections that contain densely packed neuropil (Fig. 3a). Therefore, we examined the utility of nanobodies for the labeling of endogenous synaptic molecules in 6 µm brain sections that were not subjected to any treatment except the fixation with 4% paraformaldehyde. We reasoned that thin sections might be more suitable for subsequent super-resolution imaging, where tissue depth can play a significant role [44].

**Figure 3.**
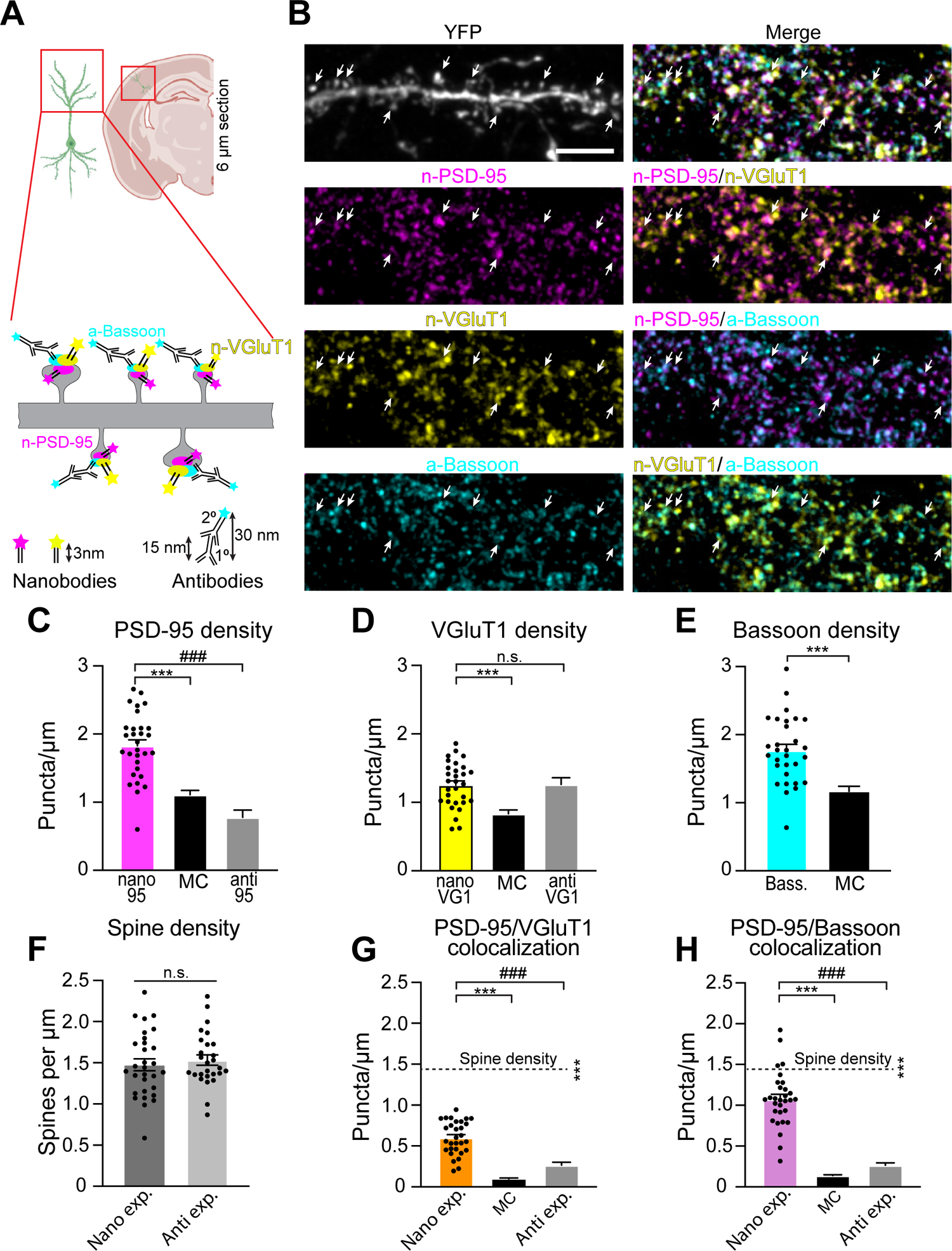
Single-domain nanobodies improve the efficiency of synaptic labeling *in situ*. **(A)** Schematic of synaptic immunolabeling with nanobodies and antibodies. **(B)** Representative four-channel confocal images of L5 apical dendrite from Thy1-YFP-H mice in 6 µm cortical cryosections. GFP antibody (gray) was used to enhance YFP fluorescence. PSD-95 (magenta) and VGluT1 (yellow) were visualized using single-domain camelid nanobodies directly conjugated to Abberior STAR 635P (n-PSD-95) and Abberior STAR 542 (n-VGluT1), respectively. Bassoon (cyan) was labeled using primary and secondary antibodies. Arrows indicate PSD-95 labeling in dendritic spines. Scale bar, 5 µm. **(C)** Comparison of average PSD-95 cluster densities per YFP labeled dendrite by nanobodies (n = 30 neurons) to Monte Carlo (MC) simulations (***p<0.0001, t-test) or antibody labeled PSD-95 cluster densities from figure 2b (^###^p<0.0001, t-test). **(D)** Comparison of average VGluT1 cluster densities per YFP labeled dendrite by nanobodies (n = 30 neurons) to MC simulations (***p<0.0001, t-test) or antibody labeled VGluT1 cluster densities from figure 2c (p = 0.9298, t-test). **(E)** Comparison of Bassoon cluster densities per YFP labeled dendrite by conventional antibodies (n = 30 neurons) to MC simulations (***p<0.0001, t-test). **(F)** Average dendritic spine densities along YFP labeled apical dendrites of L5 neurons in cryosections used for nanobody labeling (n = 30 neurons) and untreated cryosections used for antibody labeling from Fig. 2E (n = 27 neurons; p = 0.5532, t-test). **(G)** Densities of colocalized nanobody-labeled PSD-95 and VGluT1 clusters along YFP-labeled dendrites of L5 neurons compared to MC simulations (***p<0.0001, t-test) or colocalized cluster densities labeled by antibodies in untreated tissue from figure 2f (^###^p<0.0001, t-test). Dotted line indicates average spine densities as calculated in **F** (***p<0.0001, t-test). (H) Comparison of colocalized nanobody-labeled PSD-95 and antibody-labeled Bassoon cluster densities along the apical shafts of L5 neurons to MC simulations (***p<0.0001, t-test) or colocalized cluster densities labeled by antibodies in untreated tissue from Fig. 2G (^###^p<0.0001, t-test). Dotted line indicates average spine densities as calculated in **F** (***p<0.0001, t-test). Bar graphs represent the mean ± SEM. For each condition, data were collected from a minimum of ten different neurons (dots on graph) from brain sections acquired from at least three different male or female Thy1-YFP-H mice.

Cryosections from Thy-1-YFP-H animals were labeled with nanobodies for PSD-95 and VGluT1 directly conjugated to Abberior STAR 635P and Atto 542 fluorophores, respectively. We co-immunostained the same cryosections with the conventional antibody against Bassoon and determined pre- and post-synaptic puncta densities along the YFP-labeled apical dendrites of L5 pyramidal neurons (Fig. 3a, b). The PSD-95 nanobody does not recognize other MAGUK family members [52]. The specificity of the VGluT1 nanobody was examined by co-immunostaining 6 µm brain sections with VGluT1 nanobody, VGluT1 antibody and VGluT2 antibody. We observed ∼ 70% colocalization between VGluT1 antibody and nanobody, but only ∼ 20% colocalization between VGluT1 nanobody and VGluT2 antibody (Fig. S1).

PSD-95 puncta density determined by the nanobody labeling was significantly higher than MC simulations even without a need for AR or CUBIC (Fig. 3c). Importantly, PSD-95 puncta density determined by nanobody labeling nearly doubled the density of PSD-95 clusters determined by PSD-95 antibody immunostaining in untreated 6 µm brain sections (Figs. 2b and 3c). Nanobody labeling of VGluT1 in the same sections also resulted in significantly higher VGluT1 puncta density than what would be expected from random cluster distribution (Fig. 3d). However, VGluT1 puncta densities identified by nanobody labeling were indistinguishable from VGluT1 puncta density determined by antibody immunostaining in 6 µm untreated brain sections (Figs. 2c and 3d). These results indicate that conventional antibodies can efficiently label pre-synaptic specializations in thin cryosections. Consistent with this idea, immunostaining for Bassoon with the conventional antibody resulted in a significantly higher Bassoon puncta density compared to Monte Carlo simulations (Fig. 3e).

Next, we determined whether the labeling of putative synapses was also improved using nanobodies by comparing colocalized pre- and post-synaptic marker densities to dendritic spine densities. As expected, spine densities were similar to our antibody labeling experiments using 50 µm and 6 µm brain sections (Figs. 1e, 2e, and 3f). Densities of colocalized nanobody-labeled PSD-95 and VGluT1 puncta or nanobody-labeled PSD-95 and antibody-labeled Bassoon co-clusters were significantly higher than Monte Carlo simulations (Fig. 3g, h). Importantly, labeling with nanobodies significantly increased colocalized PSD-95/VGluT1 and PSD-95/Bassoon puncta densities compared to colocalized cluster densities identified by conventional antibodies in 6 µm untreated sections (Figs. 2f, g, and 3g, h). Based on our measurements of dendritic spine densities, the mean density of colocalized PSD-95 and Bassoon puncta identified ∼ 73% of putative excitatory synapses. This was a stark increase from 25% of putative synapses identified by the colocalization of PSD-95 and Bassoon antibodies (Fig. 2g). Nonetheless, the density of nanobody labeled PSD-95 clusters that colocalized with either VGluT1 or Bassoon was still significantly lower than the density of dendritic spines. Altogether, our results indicate that nanobodies significantly improve the labeling of proteins in PSDs. However, conventional antibodies efficiently label pre-synaptic specializations. Overall, nanobodies are better than conventional antibodies for quantitative measurements of putative synapses *in situ*.

### Nanobodies outperform conventional antibodies in obtaining sub-diffraction resolution of synaptic proteins using STED imaging

Molecular constituents of active zones and PSDs are diffraction-limited, which provides significant challenges for understanding the biology and function of synapses. Super-resolution imaging has provided groundbreaking discoveries in both the structure and function of the synapse, painting synapses as complex nanoscale compartments with exquisitely precise arrangements of scaffolds, receptors and neurotransmitter release sites [15]. STED microscopy enables super-resolution imaging in multiple channels with minimal chromatic aberration in XY and Z planes [18, 19]. A recent advancement in this technology, tau-STED, combines optical signals from conventional STED with physical information of the fluorescent lifetime imaging (FLIM) of individual photons [53]. Elimination of uncorrelated background noise at significantly lower excitation and STED powers increases resolution and reduces photobleaching and phototoxicity that result from repeated imaging of single regions during the Z-acquisition needed to generate three-dimensional images. Tau-STED can discriminate two objects separated only by 50 nm in the XY plane without a need for deconvolution (Fig. S2a-d) [18]. All our images are collected using Z-resolved STED, which allows for the separation of two objects in the axial plane that are only 90 nm apart (Fig. S2e-g, j).

This X, Y and Z resolution allows us to reconstruct sub-diffraction objects in three dimensions (Fig. S2h, i, k). Because of their small size and direct conjugation to fluorophores that eliminates the need for secondary antibodies, which reduces the linkage error, nanobodies have the potential to provide an increased resolution of synaptic molecules in the brain.

Indeed, nanobodies directed against GFP have been used for super-resolution imaging of overexpressed GFP-fusion proteins *in vitro* and in yeast [47]. We tested whether nanobodies enhance resolution of endogenous synaptic molecules by subjecting rat cortical neurons to STED imaging. Neurons were transfected with only GFP at the Day In Vitro (DIV) 3 to highlight neuronal morphology and visualize dendritic spines. At DIV 21, when spines are mature, we fixed these cells and labeled them with the PSD-95 nanobody directly conjugated to the Abberior STAR 635P fluorophore. We immunostained the same neurons with the antibody recognizing PSD-95, which was then subjected to indirect immunolabeling using the secondary antibody conjugated to Alexa Fluor 594. We then imaged PSD-95 puncta using conventional confocal, raw STED and tau-STED microscopy to compare the resolution of PSD-95 clusters labeled by the two reagents (Fig. 4a). As expected, raw STED significantly reduced PSD-95 cluster sizes identified by both reagents compared to confocal imaging (Fig. 4c, d).

**Figure 4.**
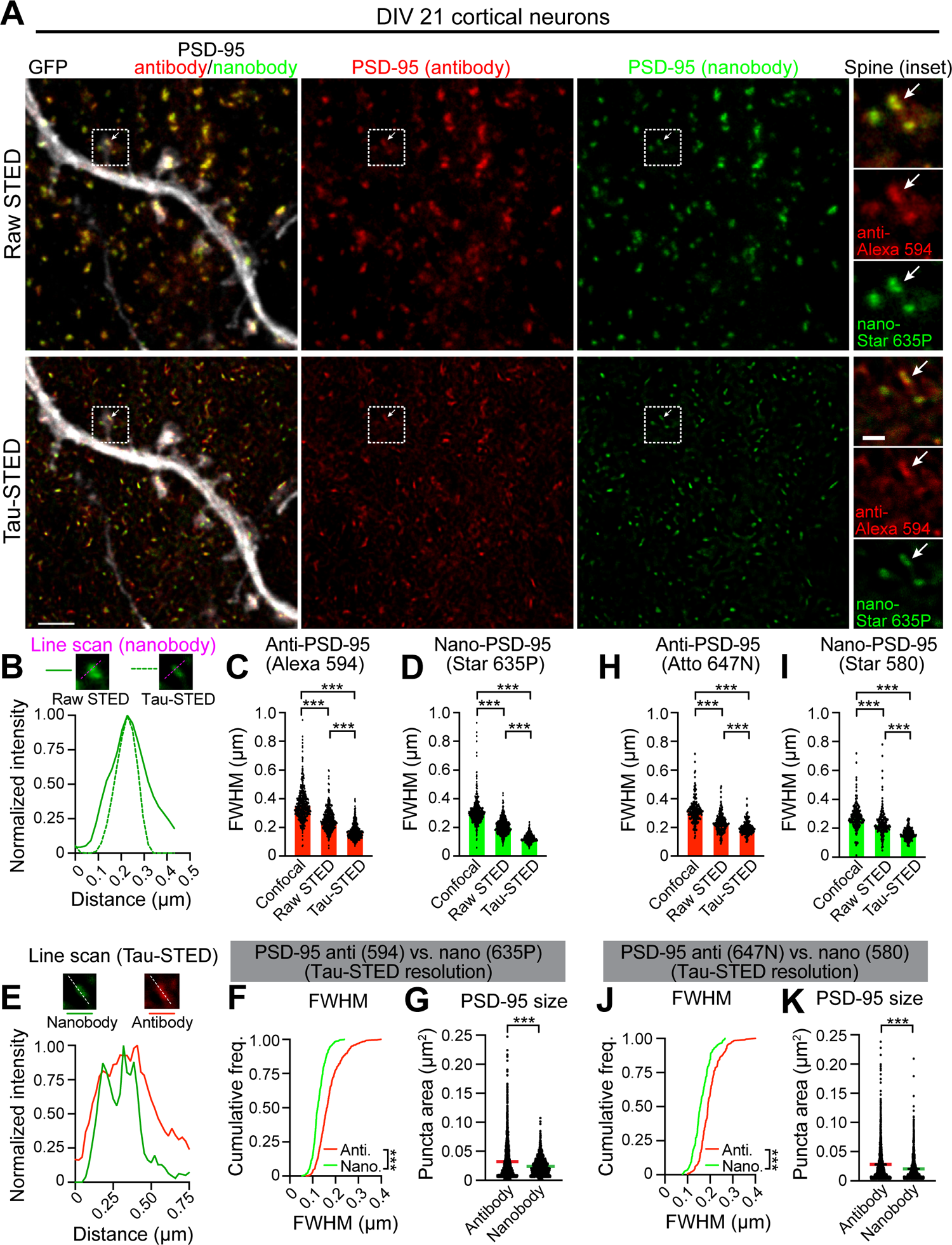
Nanobodies perform better than conventional antibodies in obtaining sub-diffraction resolution of synaptic proteins using STED imaging. **(A)** Representative raw STED and tau-STED images of the same GFP-transfected (gray) dendrite from DIV21 cortical neurons. Cultures were co-stained with the mouse PSD-95 antibody visualized using Alexa Fluor 594 conjugated secondary antibody (red) and PSD-95 nanobody directly conjugated to Abberior STAR 635P (green). The inset (square) shows a spine co-labeled with PSD-95 antibody and nanobody (arrow). Scale bars: 2 µm, inset: 500 nm. **(B)** Line profiles across a single PSD-95 nanocluster (arrows in panel **a**) in raw and tau-STED images (dotted magenta lines in insets). **(C)** Average fluorescent widths at half maxima (FWHM) of the PSD-95 antibody-labeled clusters (visualized by Alexa Fluor 594) in confocal (n = 429 clusters), raw STED (n = 444 clusters) and tau-STED modes (n = 458 clusters, ***p<0.0001, ANOVA, Tukey’s post-hoc). **(D)** Average FWHM of the PSD-95 nanobody (Aberior STAR 635P) labeled clusters in confocal (n = 425 clusters), raw STED (n = 447 clusters) and tau-STED modes (n = 469 clusters, ***p<0.0001, ANOVA, Tukey’s post-hoc). **(E)** Line profiles of a tau-STED resolved PSD-95 nanocluster (arrows in panel **a**) co-labeled with PSD-95 antibody and PSD-95 nanobody (dotted white lines in insets). **(F)** Cumulative frequency distributions of FWHM of tau-STED resolved PSD-95 nanoclusters labeled by antibody (red line, n = 458 nanoclusters) and nanobody (green line, n = 469 nanoclusters; ***p<0.0001, K-S test). **(G)** Average areas of tau-STED resolved PSD-95 nanoclusters labeled by antibody (red line, n = 2489 nanoclusters) and nanobody (green line, n = 1519 nanoclusters; ***p<0.0001, t-test). **(H)** Average FWHM of antibody-labeled PSD-95 clusters (visualized by Atto 647N) in three different imaging modes (***p<0.0001, ANOVA, Tukey’s post-hoc, confocal: n = 186, raw STED: n = 188, tau-STED: n = 189 clusters). **(I)** Average FWHM of nanobody labeled PSD-95 clusters (directly conjugated to Abberior STAR 580) in three different imaging modes (***p<0.0001, ANOVA, Tukey’s post-hoc, confocal: n = 184, raw STED: n = 188, tau-STED: n = 189 clusters). **(J)** Cumulative frequency distributions of FWHM of tau-STED resolved PSD-95 nanoclusters labeled by antibody (visualized by Atto 647N, red line, n = 189 nanoclusters) and nanobody directly conjugated to Abberior STAR 580 (green line, n = 189 nanoclusters; ***p<0.0001, K-S test). **(K)** Average areas of tau-STED resolved PSD-95 nanoclusters labeled by antibody (visualized by Atto 647N, red line, n = 1744 nanoclusters) and nanobody directly conjugated to Abberior STAR 580 (green line, n = 1441 nanoclusters; ***p<0.0001). Bar graphs represent mean ± SEM. Data was collected from a minimum of three different neurons acquired from three independent transfection experiments. Dots on bar graphs represent individual PSD-95 clusters.

Sizes of PSD-95 antibody (Fluorescence Width at Half Maximum – FWHM = 255 ± 3.6 nm) and PSD-95 nanobody (FWHM = 209 nm ± 2.5 nm) labeled puncta imaged by raw STED were similar to previously reported values [19, 20]. Tau-STED further reduced sizes of PSD-95 nanoclusters identified by both PSD-95 antibody (FWHM = 172 ± 2.4 nm) and PSD-95 nanobody (FWHM = 126 ± 1.3 nm, dotted magenta line in Fig. 4b) compared to the raw STED regiment without the need for deconvolution (Fig. 4b-d).

Notably, the same PSD-95 nanoclusters were always better resolved in the nanobody channel than the antibody channel. The line plot of the same PSD-95 nanocluster imaged in the tau-STED mode (Fig. 4e, dotted white line) revealed two peaks spaced ∼ 130 nm apart for PSD-95 nanobody. In contrast, the same cluster labeled by the antibody showed only a single peak (Fig. 4e). Cumulative probability distributions of FWHM and average PSD-95 nanocluster areas further demonstrate that PSD-95 nanoclusters are significantly smaller when imaged with nanobodies (0.023 ± 0.0003 µm^2^) than with antibodies (0.032 ± 0.0006 µm^2^; Fig. 4f, g). We obtained similar results when we switched fluorophores, using Abberior STAR-580 conjugated PSD-95 nanobody and Atto 647N conjugated secondary antibody to amplify the PSD-95 primary antibody signal (Fig. 4h-k). Thus, the improved resolution of PSD-95 nanoclusters provided by the nanobody labeling is not due to the chromatic aberration or the mix of secondary antibodies used in our labeling protocol. These results indicate that nanobodies outperform conventional antibodies in obtaining sub-diffraction resolution enabled by FLIM-based tau-STED super-resolution microscopy.

### Nanobody labeling enhances the localization of PSD-95 to sites of neurotransmitter release in brain sections

Previous work used two-color STED nanoscopy to determine the nano-organization of VGluT1 and PSD-95 labeled by antibodies in dendritic spines of thick (∼ 250 µm) brain sections that were cleared with CUBIC to minimize light scattering that limits STED resolution in tissue [19]. However, the necessity of the refractive index matching of CUBIC cleared sections and the loss of resolution that occurs with increasing depth of the section beyond a few microns limits the accurate localization of proteins to active zones and PSDs. Therefore, we reasoned that combining tau-STED nanoscopy with nanobody labeling of thin (6 µm) brain sections might improve the quality of images and increase the accuracy of synaptic protein localization to trans-synaptic nanodomains *in situ*.

We collected 6 µm cryosections from Thy1-YFP-H brains and immunolabeled them with GFP antibody to enhance the YFP signal and identify dendritic spines. Based on our confocal data demonstrating that nanobodies and antibodies are equally effective in labeling pre-synaptic specializations (Fig. 3), we used the Bassoon antibody to visualize active zones. We then used either PSD-95 nanobody or the conventional PSD-95 antibody to label post-synaptic specializations. We imaged YFP in the confocal mode. PSD-95 and Bassoon were imaged using a 3D resolved two-color tau-STED regime to visualize their nano-organization in YFP-labeled spines (Fig. 5a-f). To confirm the correct localization of STED-resolved pre- and post-synaptic nanoclusters to spines and each other *in situ*, we 3D-rendered confocal and tau-STED images in Neurolucida 360 (Fig. 5e, f). These 3D tau-STED experiments revealed that nanobody-resolved PSD-95 nanoclusters in spines were directly juxtaposed to Bassoon nanoclusters (Fig. 5e). In contrast, PSD-95 antibody poorly identified trans-synaptic nanodomains in spines and Bassoon nanoclusters were often found without any aligned PSD-95 nanoclusters (Fig. 5f).

**Figure 5.**
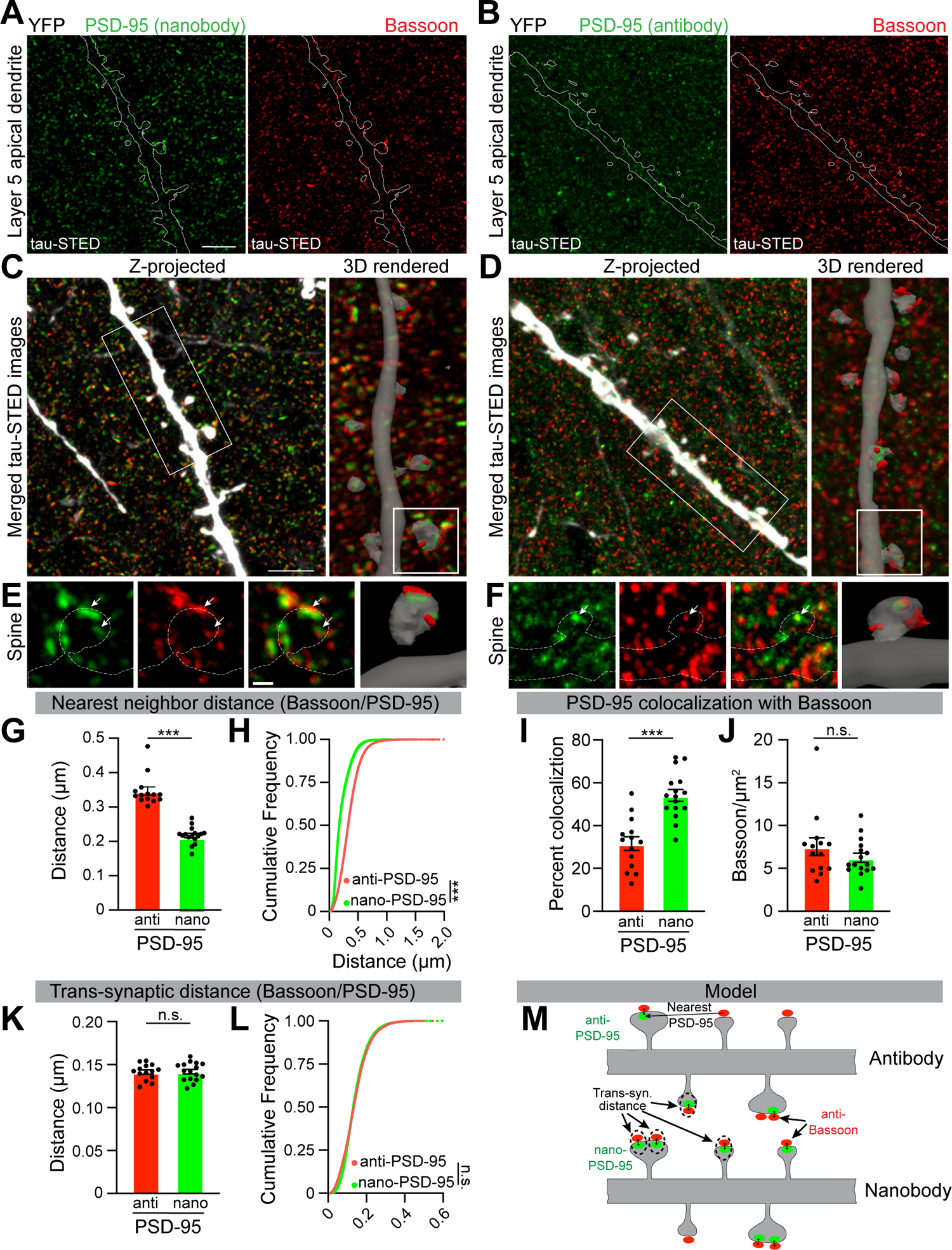
PSD-95 nanobody exhibits significantly better nano-localization to active zones than PSD-95 antibody. **(A, B)** Representative two-channel tau-STED images of PSD-95 (green) labeled by nanobody (**a**) or antibody (**b**) and Bassoon (red) along the apical dendrite of an L5 pyramidal neuron in Thy-1-YFP-H mice. YFP (dotted outline, enhanced by labeling with GFP antibody) was imaged in confocal resolution. Scale bar, 3 µm. **(C, D)** Maximum projection (left) and 3D rendering (right, inset) of merged images shown in panels **A, B**. Scale bar, 3 µm. **(E, F)** Dendritic spines (dotted white outlines) from the reconstructed dendritic segments (rectangular insets in **C, D**) containing PSD-95 (green) nanoclusters aligned to Bassoon (red) nanoclusters (arrows). Spines were reconstructed in 3D using Neurolucida 360 to verify the localization of PSD-95 and Bassoon nanoclusters to the YFP-labeled spine. Scale bar, 500 nm. **(G)** Average center-to-center distances between Bassoon and the nearest PSD-95 nanoclusters in a 25 x 25 µm imaging field identified by staining for antibody (n = 14 fields) or nanobody (n = 16 fields, ***p<0.0001, t-test). **(H)** Cumulative frequency distribution of the nearest neighbor distances between centers of individual Bassoon and PSD-95 nanoclusters identified either by staining for PSD-95 antibody (n = 68806 clusters) or PSD-95 nanobody (n = 39405 clusters, ***p<0.0001, K-S test). **(I)** Average colocalization probability between Bassoon and PSD-95 nanoclusters in a 25 x 25 µm imaging field determined by staining for PSD-95 antibody (n = 14 fields) and nanobody (n = 16 fields, ***p<0.0001, t-test). **(J)** Average Bassoon nanocluster density in 25 x 25 µm images stained with PSD-95 antibody (n = 14 images) and nanobody (n = 16 images, p = 0.2518, t-test). **(K)** Average center-to-center distances between colocalized Bassoon and PSD-95 nanoclusters labeled by PSD-95 antibody (n = 14 images) and nanobody (n = 16 images, p = 0.9037, t-test). **(L)** Cumulative frequency distributions of center-to-center distances between colocalized Bassoon and PSD-95 nanoclusters labeled by PSD-95 antibody (n = 12992 clusters) and nanobody (n = 26092 clusters, p=0.4483, K-S test). **(M)** Model representing synaptic PSD-95 localization identified either by antibody or nanobody to active zones marked by Bassoon.

We quantified the degree of PSD-95 juxtaposition to neurotransmitter release sites by measuring the nearest neighbor distances between centers of Bassoon and PSD-95 nanoclusters visualized either by the PSD-95 nanobody or the PSD-95 antibody. Segmentation and subsequent analysis of X, Y and Z-STED-resolved Bassoon and PSD-95 nanoclusters were performed in an unbiased manner using the DiAna macro in ImageJ [18, 54]. Bassoon’s nearest PSD-95 neighbor was, on average, within 200 nm when we used the PSD-95 nanobody (Fig. 5g). However, the average distance to the nearest active zone was significantly further (350 nm) when PSD-95 nanoclusters were labeled with the antibody. Indeed, the entire cumulative distribution of center-to-center distances between Bassoon and PSD-95 was shifted to the right when we used the PSD-95 antibody, indicating greater distances between the individual PSD-95 and Bassoon nanoclusters (Fig. 5h). These data indicate that the PSD-95 nanobody significantly increases the accuracy of post-synaptic nanocluster localization opposite of active zones in brain sections. Consistent with these results, ∼60% of PSD-95 nanoclusters labeled by the nanobody localized near Bassoon nanoclusters compared to only 30% of antibody-labeled PSD-95 nanoclusters (Fig. 5i). This difference was due poor efficiency of antibody labeling of endogenous PSD-95, because the density of endogenous Bassoon nanoclusters was similar in both conditions (Fig. 5j).

Next, we analyzed the center-to-center distances of PSD-95/Bassoon nanoclusters that colocalized at the nanoscale and thus were deemed trans-synaptic contacts. Using these parameters, average distances and cumulative probability distributions between centers of Bassoon and PSD-95 nanoclusters identified either by the PSD-95 antibody or PSD-95 nanobody were nearly identical in tau-STED (Fig. 5k, l; antibody, d=141.4 nm ± 0.0025 nm; nanobody, d=141.8 nm ± 0.0026 nm). These data indicate that both PSD-95 antibody and PSD-95 nanobody can localize PSD-95 clusters near active zones marked by Bassoon. However, PSD-95 antibody identifies only a subset (∼30%) of trans-synaptic nanomodules, which could have negative consequences for the quantification of synaptic nano-architecture in the brain tissue (Fig. 5m). Thus, labeling with PSD-95 nanobody increases the accuracy of trans-synaptic nano-organization in brain sections.

### TC synapses in layer 5A of S1 are larger than neighboring CC synapses despite similar trans-synaptic nano-organization

Having characterized tools for STED imaging *in situ*, we next sought to determine how functionally diverse excitatory synapses in the brain organize their nano-architecture. L5 pyramidal neurons are a great model for examining this question due to their vast dendritic trees spanning layers 1 to 5. In S1, they receive glutamatergic input from CC and TC afferents in both apical and basal dendritic domains that have been functionally characterized *in vivo* [35, 55]. In the brain, CC and TC synapses can be distinguished by the presence of either vesicular glutamate transporter 1 (VGluT1) or VGluT2, respectively, allowing for the direct comparison of these glutamatergic synaptic sub-types in the same cell [56, 57].

We initially focused on the nano-organization of TC synapses on L5A basal dendrites where afferents from the posterior-medial (POm) thalamic nucleus generate robust post-synaptic responses [55]. We collected 6 µm brain sections from S1 of P35 – P90 Thy1-YFP-H mice in which L5 pyramidal neurons genetically express high levels of YFP [50]. We immunolabeled these sections with the nanobody against VGluT1 (a marker of CC synapses), the antibody against VGluT2 (a marker of TC synapses), the antibody against Bassoon (active zones), and the nanobody against PSD-95 (PSDs).

We then performed simultaneous five-channel confocal and STED imaging (Fig. 6a). YFP (enhanced by GFP immunostaining), VGluT1 and VGluT2 were imaged in confocal resolution to visualize dendritic spines on L5 pyramidal cells and to distinguish between TC and CC synapses. PSD-95 and Bassoon were imaged in Z-resolved (3D) tau-STED to determine the trans-synaptic nano-organization in VGluT1 and VGluT2 spines. We acquired ∼2 µm stacks through basal dendrites in L5A. Both confocal and STED images were reconstructed in Neurolucida 360 to obtain a 3D rendering of dendritic spines and associated synaptic clusters (Fig. 6a, b). After 3D reconstructions, only those synaptic clusters that colocalized with YFP-labeled spines were analyzed (Fig. 6a, top right). This approach allowed us to identify spines receiving either VGluT1 or VGluT2 inputs (Fig. 6b, top row). We visualized trans-synaptic nano-organization by localizing PSD-95 and Bassoon nanoclusters to spines belonging to either category (Fig. 6b, middle and bottom rows). Due to the high complexity of spines and synaptic clusters in a stack, we only show the rendering of sub-stacks through individual spines. Maximum projections showing all spines and associated clusters in each 2 µm stack are shown in Fig. S3.

**Figure 6.**
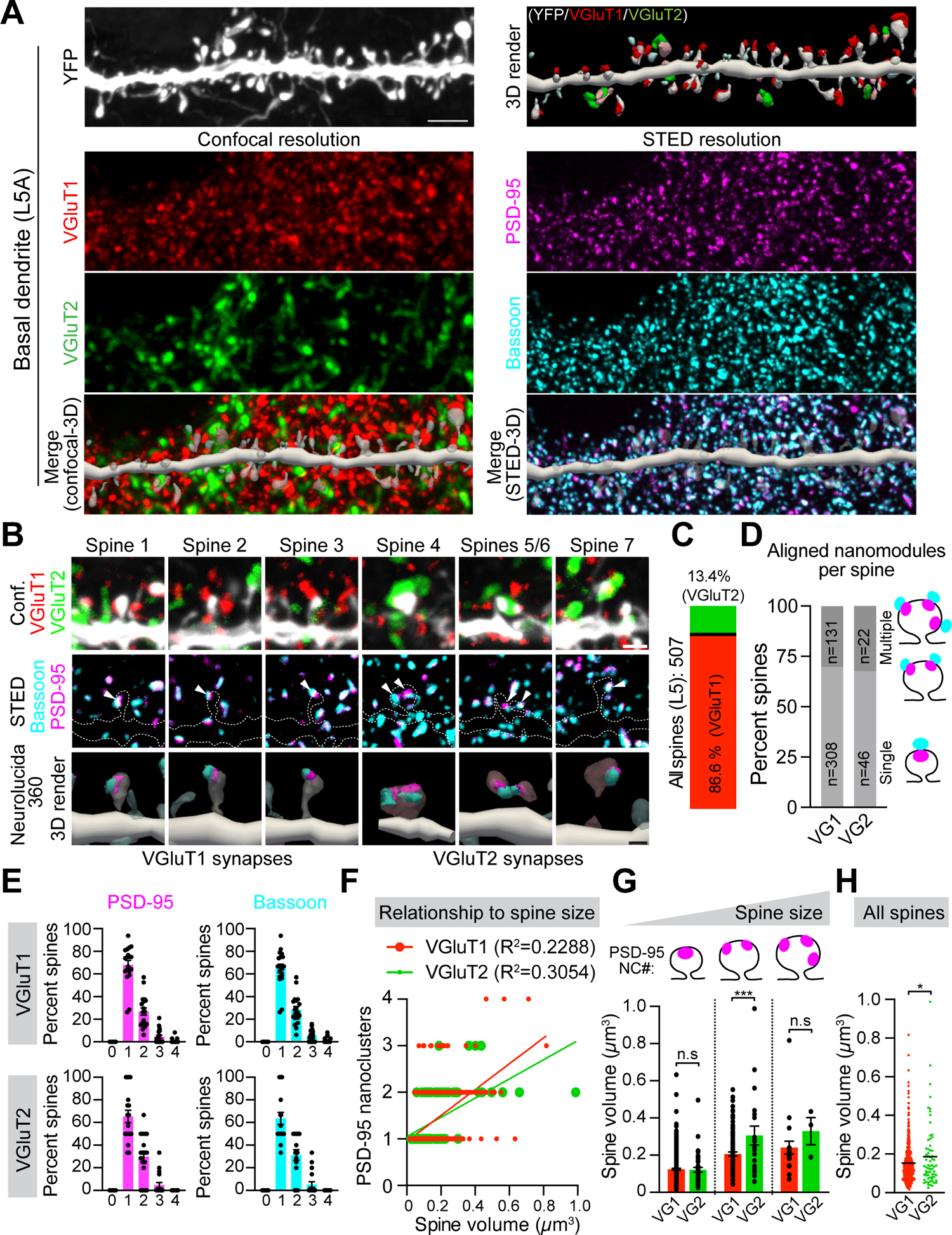
TC synapses in layer 5A of S1 are larger than neighboring CC synapses despite similar trans-synaptic nano-organization. **(A)** Representative 5 channel confocal/tau-STED images of YFP labeled dendritic spines on basal dendrites of genetically identified L5 pyramidal neurons in S1 of Thy-1-YFP-H mice. YFP (white, enhanced by the labeling with GFP antibody), VGluT1 (red, nanobody/Atto 542) and VGluT2 (green, Alexa Fluor 790) were imaged in confocal resolution (left panel). PSD-95 (magenta, nanobody/Abberior STAR 635P) and Bassoon (cyan, antibody/Alexa Fluor 594) were imaged in STED resolution (right panel). Confocal and STED channels were reconstructed in 3D using Neurolucida 360 to categorize spines as VGluT1^+^ or VGluT2^+^ and verify the localization of PSD-95 and Bassoon nanoclusters to individual spines. Scale bar, 2 µm. **(B)** Representative sub-stacks of individual dendritic spines that received either VGluT1 (red) or VGluT2 (green) glutamatergic input. The top row shows merged confocal images of either VGluT1^+^ or VGluT2^+^ YFP-labeled spines. The middle row shows tau-STED-resolved PSD-95 (magenta) and Bassoon (cyan) nanoclusters (arrows) in the same spines (dotted outlines). The bottom row shows Neurolucida 360 3D-reconstructed spines and corresponding PSD-95 and Bassoon nanoclusters. High-contrast images are shown. Scale bars, 1 µm (tau-STED), 500 nm (3D render). **(C)** Quantification of the fraction of TC and CC synapses determined by VGluT1 and VGluT2 staining of 507 spines from 19 different dendritic segments. **(D)** Quantification of the fraction of VGluT1^+^ (n = 439) and VGluT2^+^ (n = 68) spines with single or multiple (two or more) aligned PSD-95 and Bassoon nanoclusters. **(E)** Percentage of VGluT1^+^ (n = 439) and VGluT2^+^ (n = 68) spines containing one, two, three and four PSD-95 and Bassoon clusters from 19 dendritic segments. **(F)** Quantification of the relationship between spine size and numbers of PSD-95 nanoclusters in VGluT1^+^ (R^2^ = 0.2288, slope = 2.781 ± 0.48) and VGluT2^+^ (R^2^ = 0.3054, slope = 1.942 ± 0.72) spines (p = 0.0711, ANCOVA). **(G)** Comparison of PSD-95 relationship to spine size in VGluT1^+^ and VGluT2^+^ spines with single (VG1: n = 308 spines, VG2: n = 44 spines, p = 0.3484, t-test), two (VG1: n = 107 spines, VG2: n = 21 spines, ***p = 0.0008, t-test) and three (VG1: n = 20 spines, VG2: n = 3 spines, p = 0.3582, t-test) PSD-95 nanoclusters (NC). **(H)** Average sizes of all VGluT1^+^ (n = 438) and VGluT2^+^ (n = 68) spines regardless of the number of nanoclusters (*p = 0.0263, t-test). Bars represent the mean ± SEM acquired from spines of 19 different basal dendritic segments of L5 neurons obtained from two biological replicates.

We imaged a total of 507 dendritic spines on basal dendrites. Following dendritic segment reconstructions, only 68 (13.4%) spines were determined to be TC synapses based on VGluT2 colocalization. The remaining 439 (86.6%) spines contained VGluT1 immunolabeling and thus were identified as CC synapses (Fig. 6c). A small number of spines could not be fully reconstructed or reliably assigned to either VGluT1 or VGluT2 category due to occlusion by other spines, dendrites or axons in the image field. These spines were not included in the analysis. Following the identification of CC or TC synapses, we next determined the nano-organization of PSD-95 and Bassoon nanoclusters in each spine (Fig. 6d). Discrete, aligned PSD-95 and Bassoon nanoclusters were identified as trans-synaptic nanomodules [18, 19]. Consistent with the published literature, the majority (∼70%) of VGluT1^+^ spines contained a single aligned PSD-95/Bassoon nanomodule and ∼30% of VGluT1^+^ spines contained multiple (two, three or four) aligned nanomodules (Fig. 6b, d). Similar to VGluT1^+^ spines, 68% of VGluT2^+^ spines had only one aligned PSD-95/Bassoon nanomodule and 32% had multiple aligned nanomodules (Fig. 6b, d). Moreover, distributions of individual PSD-95 and Bassoon nanoclusters in VGluT1^+^ and VGluT2^+^ spines were similar (Fig. 6e). Thus, the highest proportion of spines in both glutamatergic synapse categories contained only one PSD-95 and Bassoon nanocluster, while spines with two, three and four nanoclusters were present with a progressively lower probability.

Previous studies demonstrated that the number of PSD-95 and Bassoon nanoclusters scale with increasing spine size, consistent with the idea that larger spines contain multiple nanomodules [18, 19]. Therefore, we asked whether the same rules apply to VGluT2^+^ spines in L5A. The number of PSD-95 nanoclusters scaled linearly with spine size in both VGluT1^+^ (R^2^ = 0.2288, p<0.0001) and VGluT2^+^ spines (R^2^ = 0.3054, p<0.0001), and the relationship between the number of PSD-95 nanoclusters and spine size was similar (Fig. 6f, p = 0.0711, ANCOVA). Thus, large VGluT1^+^ and VGluT2^+^ spines tend to have multiple nanoclusters. Despite these similarities, VGluT2^+^ containing spines were larger on average than VGluT1^+^ spines (Fig. 6h), consistent with previous studies [56, 58, 59]. To determine the nanoscale logic for the overall larger size of spines receiving TC input, we compared sizes of VGluT1^+^ and VGluT2^+^ spines with one, two and three PSD-95 nanoclusters (Fig. 6g). VGluT1^+^ and VGluT2^+^ spines containing only one PSD-95 had statistically indistinguishable sizes. However, VGluT2^+^ spines containing two PSD-95 nanoclusters were significantly larger than VGluT1^+^ spines containing the same number of PSD-95 nanoclusters (Fig. 6g). We saw a similar trend for spines with three PSD-95 nanomodules; this difference was not statistically significant (p = 0.2408, t-test), likely due to a very low abundance of VGluT2^+^ spines with three nanomodules (n = 3 spines). These observations are consistent with the notion that VGluT2^+^ spines need to reach much larger sizes than neighboring VGluT1^+^ spines to adopt a multiple nanocluster architecture. Thus, unless synaptic nanodomains in VGluT2^+^ spines occupy a much lower fraction of the post-synaptic membrane than in VGluT1^+^ spines, these data raise a question about the relationship between sizes of individual PSD-95 nanoclusters and VGluT2^+^ spine volume [19].

### TC synapses in L1-3 apical tufts of L5 pyramidal cells form spines with multiple trans-synaptic nanomodules with high probability

Layer 5 pyramidal cells also receive the POm TC input within their apical domains (L1-3), which is stronger than expected from its distant localization relative to the cell body [35, 55]. Therefore, we sought to determine the nano-architecture of VGluT2^+^ containing spines on apical dendrites of L5 pyramidal neurons. We subjected the same brain sections that were used for imaging of spines on basal dendrites to five-channel confocal and tau-STED imaging of TC and CC synapses in apical tufts of L5 pyramidal neurons (Fig. 7a). We imaged YFP (enhanced by GFP immunostaining), VGluT1 and VGluT2 in confocal resolution to visualize dendritic spines and classify CC and TC synapses, respectively. PSD-95 and Bassoon were imaged in the Z-resolved tau-STED mode to identify trans-synaptic nano-organization in VGluT1^+^ and VGluT2^+^ spines. Both confocal and STED images were reconstructed in Neurolucida 360 to obtain 3D rendering of individual dendritic spines with associated synaptic clusters (Fig. 7b-d and Supplementary Video 1-3). Spines belonging to the VGluT1 or VGluT2 category were subjected to the analysis of synaptic nano-organization using tau-STED resolved PSD-95 and Bassoon nanoclusters (Fig. 7e and Fig. S4).

**Figure 7.**
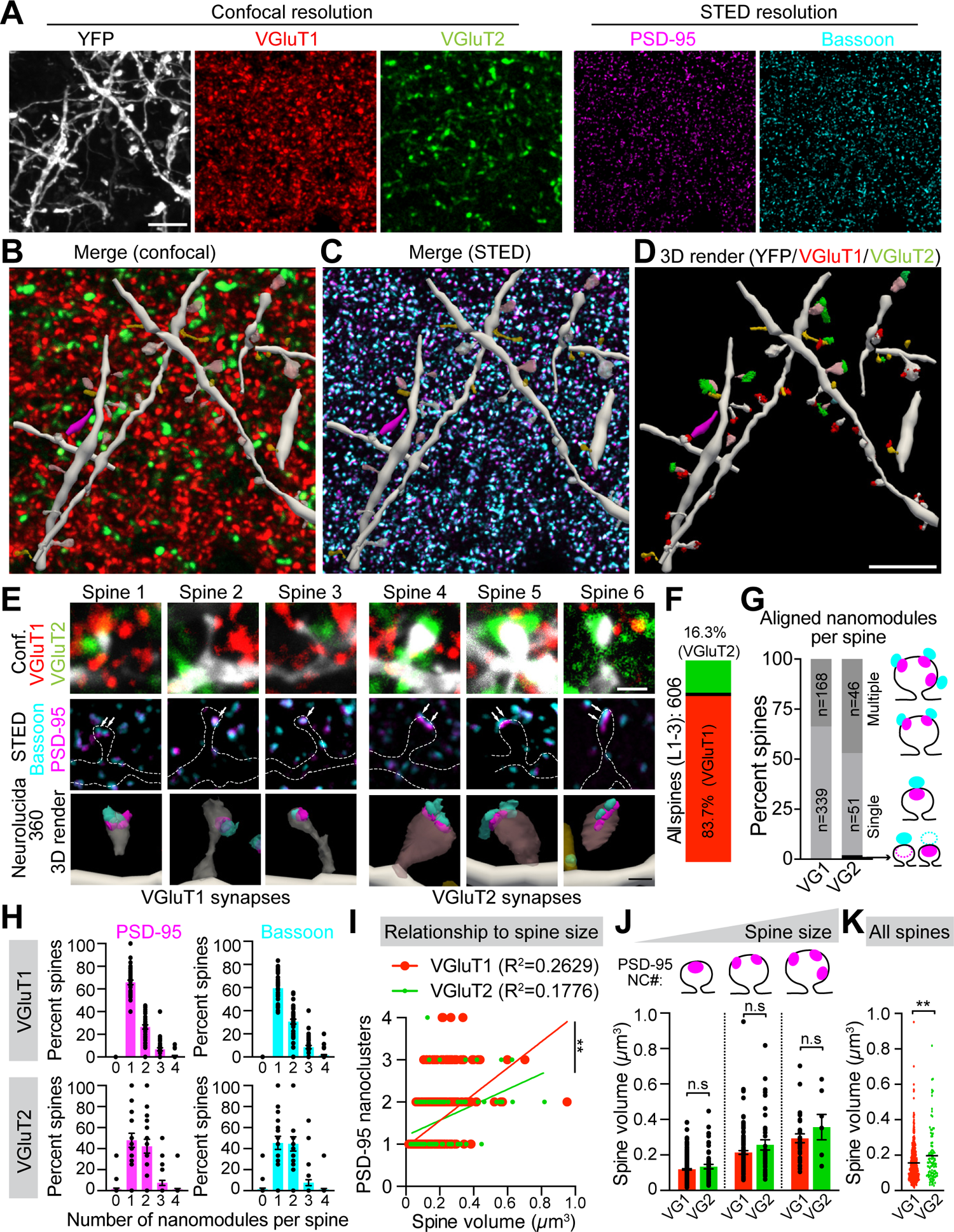
TC synapses in L1-3 apical tufts of L5 pyramidal cells form spines with multiple trans-synaptic nanomodules with high probability. **(A)** Representative 5 channel confocal/tau-STED images of YFP labeled dendritic spines on apical dendrites of genetically identified L5 pyramidal neurons from S1 of Thy-1-YFP-H mice. YFP (white, enhanced by GFP antibody), VGluT1 (red, nanobody/Atto 542) and VGluT2 (green, antibody/Alexa Fluor 790) were imaged in confocal resolution (left panels). PSD-95 (magenta, nanobody/Abberior STAR 635P) and Bassoon (cyan, antibody/Alexa Fluor 594) were imaged in tau-STED resolution (right panels). Scale bar, 5 µm. **(B-D)** Confocal and tau-STED channels were reconstructed in 3D using Neurolucida 360 to categorize spines as VGluT1^+^ or VGluT2^+^ and verify the localization of PSD-95 and Bassoon nanoclusters to individual spines. Scale bar, 5 µm. **(E)** Representative sub-stacks of individual dendritic spines that received either VGluT1 (red) or VGluT2 (green) glutamatergic input. The top row shows merged confocal images of either VGluT1^+^ or VGluT2^+^ YFP-labeled spines. The middle row shows tau-STED-resolved aligned PSD-95 (magenta) and Bassoon (cyan) nanoclusters (arrows) in the same spines (dotted outlines). The bottom row shows Neurolucida 360 3D-reconstructed spines and corresponding PSD-95 and Bassoon nanoclusters. High-contrast images are shown. Scale bars, 1 µm (tau-STED), 500 nm (3D render). **(F)** Quantification of the fraction of TC and CC synapses determined by VGluT1 and VGluT2 staining of 606 spines from L1-3 apical dendrites. **(G)** Quantification of the fraction of VGluT1^+^ (n = 507) and VGluT2^+^ (n = 99) spines with single or multiple (two or more) aligned PSD-95 and Bassoon nanomodules. **(H)** Distributions of PSD-95 and Bassoon nanocluster numbers per individual VGluT1^+^ (n = 507) or VGluT2^+^ (n = 99) spines from 37 independently imaged dendrites. **(I)** Quantification of the relationship between spine size and PSD-95 nanocluster numbers in VGluT1^+^ (R^2^ = 0.2629, slope = 3.122 ± 0.464) and VGluT2^+^ (R^2^ = 0.1776, slope = 1.833 ± 0.799) containing spines (**p = 0.0037, ANCOVA). **(J)** Average sizes of VGluT1^+^ and VGluT2^+^ spines with single (VG1: n = 329 spines, VG2: n = 49 spines, p = 0.5722, t-test), two (VG1: n = 133 spines, VG2: n = 39 spines, p = 0.0931, t-test) and three (VG1: n = 31 spines, VG2: n = 8 spines, p = 0.1368, t-test) PSD-95 nanoclusters. **(K)** Average sizes of all VGluT1^+^ and VGluT2^+^ spines regardless of the number of nanoclusters (**p = 0.0014, Student’s t-test). Bars represent the mean ± SEM acquired from spines on 37 apical dendritic segments of L5 neurons from two biological replicates.

We imaged a total of 606 dendritic spines on apical dendrites. Only 99 (16.3%) spines were determined to be TC synapses based on VGluT2 colocalization after reconstructions of individual spines. The remaining 507 (83.7%) spines were VGluT1^+^ and we classified them as CC synapses (Fig. 7f). We next determined the nano-organization of aligned PSD-95 and Bassoon nanoclusters in VGluT1^+^ and VGluT2^+^ spines (Fig. 7g). Similar to our analysis of VGluT1^+^ spines on basal dendrites of L5 neurons, 66% of VGluT1^+^ spines in L1-3 had one aligned PSD-95/Bassoon nanomodule and ∼33% of spines contained multiple (two or more) aligned PSD-95/Bassoon nanomodules. In contrast, fewer VGluT2^+^ spines (51%) contained a single aligned PSD-95/Bassoon nanomodule. Nearly half (46%) of VGluT2^+^ spines in apical tufts of L5 neurons contained synapses with multiple aligned PSD-95/Bassoon nanomodules.

These results indicate that VGluT2^+^ spines on apical dendrites of L5 neurons in S1 form synapses with multiple nanomodules more often than neighboring VGluT1^+^ spines or VGluT2^+^ spines on basal dendrites of L5 neurons (Fig. 6). To examine the nature of TC vs. CC synaptic complexity more closely, we compared the distribution of VGluT1^+^ and VGluT2^+^ spines with one, two, three or four PSD-95 and Bassoon nanoclusters (Fig. 7h). As previously established, VGluT1^+^ spines with one PSD-95 and Bassoon nanocluster were most abundant and represented ∼ 66% of all VGluT1^+^ spines examined. Only ∼25% of all VGluT1^+^ spines contained two PSD-95 and Bassoon nanoclusters. CC synapses with three and four PSD-95 and Bassoon nanoclusters collectively represented less than 10% of all VGluT1^+^ spines (Fig. 7h) [18, 19]. This distribution was shifted for VGluT2^+^ spines in a manner that led to nearly an equal proportion of spines with one and two PSD-95 and Bassoon nanoclusters. VGluT2^+^ spines with three and four nanoclusters were uncommon (< 6%), which was similar to the frequency seen for VGluT1^+^ spines (Fig. 7h). It is important to note that we saw only one or two VGluT2^+^ spines with any given nano-organization per dendritic segment due to their low abundance in the cortex and a high zoom factor necessary to obtain an appropriate pixel size for STED imaging. Thus, even though proportionally there were more VGluT2^+^ spines with multiple aligned PSD-95/Bassoon nanomodules in L1-3, VGluT1^+^ spines with two or more nanomodules were still more abundant on apical dendrites of L5 cells. Therefore, VGluT2^+^ spines with three nanomodules were extremely rare and we seldom saw a VGluT2^+^ spine with four PSD-95 nanoclusters aligned to four Bassoon nanoclusters.

Because of these differences in the pattern of PSD-95 and Bassoon nano-organization in VGluT1^+^ and VGluT2^+^ spines, we wondered whether the relationship between nanomodule numbers and spine size might also differ between TC and CC synapses in L1-3. The sizes of both VGluT1^+^ and VGluT2^+^ spines were positively correlated with PSD-95 nanocluster numbers (VGluT1: R^2^ = 0.2629, p<0.0001; VGluT2: R^2^ = 0.1776, p<0.0001, Fig. 7i). However, the slope of this linear relationship was significantly flatter for VGluT2^+^ spines (slope = 1.833, 95% confidence 1.034 to 2.633) compared to VGluT1^+^ spines (slope = 3.122, 95% confidence 2.658 to 3.586; p = 0.0037, ANCOVA). We reasoned that this is due to fewer VGluT2^+^ spines with one PSD-95 nanocluster and more VGluT2^+^ spines with two PSD-95 nanoclusters compared to VGluT1^+^ synapses. Additionally, there were fewer VGluT2^+^ spines with three or four PSD-95 nanoclusters in L1-3. To shed light on this skewed relationship, we compared sizes of VGluT1^+^ and VGluT2^+^ spines with one, two and three PSD-95 nanoclusters (Fig. 7j). There were no significant differences in average volumes of VGluT1^+^ and VGluT2^+^ spines within each PSD-95 nanocluster number category, indicating that apical VGluT1^+^ and VGluT2^+^ spines with equivalent numbers of PSD-95 nanoclusters are remarkably similar with respect to their size. Yet, when comparing all spines in L1-3 regardless of their PSD-95 nanocluster numbers, VGluT2^+^ spines were significantly larger on average than VGluT1^+^ spines (Fig. 7k). These data support our observation that VGluT2^+^ spines with multiple PSD-95/Bassoon nanomodules, which are larger than spines with one nanomodule, are common in apical domains of L5 neurons. Altogether, we show that CC and TC synapses in apical tufts of L5 neurons are very similar to each other, with the main difference being a high relative abundance of VGluT2^+^ spines with multiple pre- and post-synaptic nanomodules that results in larger average volumes of spines receiving TC input.

### TC synapses in apical and basal compartments of L5 neurons are built on distinct nanoscale principles

Our findings indicate that TC synapses in both basal and apical domains of L5 pyramidal neurons are, on average, larger than CC synapses. The larger average size of apical VGluT2^+^ spines is due to the high relative abundance of spines containing multiple nanomodules. Otherwise, apical VGluT2^+^ spines are structurally similar to VGluT1^+^ spines containing equivalent numbers of nanomodules. The same logic cannot be applied to VGluT2^+^ spines on basal dendrites. Indeed, most basal VGluT2^+^ spines contain a single aligned PSD-95/Bassoon nanocluster. Nonetheless, basal VGluT2^+^ spines are still larger than neighboring VGluT1^+^ spines. These data imply that TC synapses in apical and basal domains might follow distinct nanoscale principles of organization.

To begin to address this question, we analyzed sizes of individual PSD-95 nanoclusters in VGluT2^+^ spines from L5A and L1-3 (Fig. 8). Our tau-STED imaging indicates that individual PSD-95 nanoclusters in VGluT2^+^ spines on basal dendrites are significantly larger on average than PSD-95 nanoclusters in VGluT2^+^ spines on apical dendrites of L5 neurons (Fig. 8a, b). Thus, VGluT2^+^ spines in L5A and L1-3 use nanoclusters of different sizes to build their nano-architecture. These data raise a question of whether VGluT2^+^ spines in either dendritic domain conform to modular rules of assembly where the number but not the size of PSD-95 nanoclusters scales with spine size, as previously demonstrated for VGluT1 synapses [19]. Therefore, we measured the sizes of individual PSD-95 nanoclusters in apical (L1-3) and basal (L5A) VGluT2^+^ spines containing one, two and three nanoclusters. Volumes of individual PSD-95 nanoclusters in VGluT2^+^ spines in L1-3 did not significantly differ between spines with different numbers of PSD-95 nanoclusters (Fig. 8c). These findings are consistent with modular principles of nano-organization. In contrast, the sizes of individual PSD-95 nanoclusters in VGluT2^+^ spines on basal dendrites were more variable. Here, VGluT2^+^ spines with one PSD-95 had significantly smaller PSD-95 nanoclusters than VGluT2^+^ spines with two PSD-95 (Fig. 8d). Interestingly, sizes of individual PSD-95 nanoclusters in VGluT2^+^ spines with two and three PSD-95 were similar (Fig. 8d). This effect could reflect capping of individual nanocluster sizes in spines with multiple (two or more) nanoclusters. Whether nanocluster sizes are restricted by some mechanism or continue to increase as spines grow is difficult to assess from current data due to a very low number (n = 3) of basal VGluT2^+^ spines with three PSD-95 nanodomains. Because nanocluster sizes vary between spines containing one and two PSD-95 nanoclusters, these findings suggest that VGluT2^+^ spines on basal dendrites may not follow modular principles of nano-organization.

**Figure 8.**
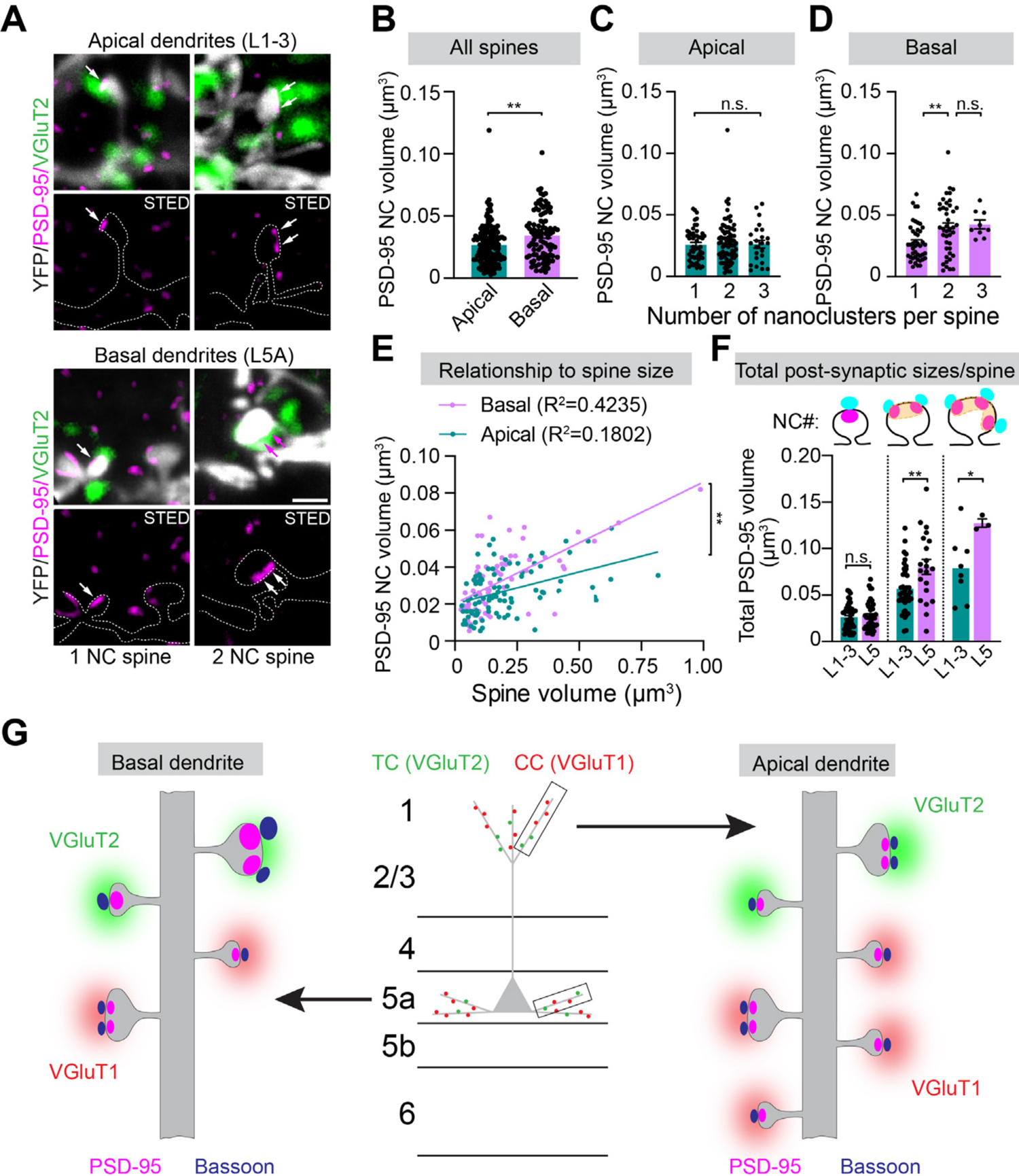
TC synapses in apical and basal compartments of L5 neurons are built on distinct nanoscale principles. **(A)** Representative images of single and two PSD-95 nanocluster (magenta, arrows) VGluT2^+^ (green) YFP labeled spines (gray, dotted outlines) from apical and basal dendritic compartments of L5 pyramidal neurons in S1 of Thy-1-YFP-H mice. Scale bar, 1 µm. **(B)** Average PSD-95 nanocluster (NC) volumes in all VGluT2^+^ spines from apical (n = 152 nanoclusters) and basal (n = 95 nanoclusters) dendrites (**p = 0.0011, Student’s t-test). **(C)** Average PSD-95 nanocluster (NC) volumes in VGluT2^+^ spines with one (n = 52 nanoclusters), two (n = 76 nanoclusters) and three (n = 24 nanoclusters) PSD-95 on apical dendrites (p = 0.6376, ANOVA). **(D)** Average PSD-95 nanocluster (NC) volumes in VGluT2^+^ spines with one (n = 44 nanoclusters), two (n = 42 nanoclusters) and three (n = 9 nanoclusters) PSD-95 on basal dendrites (**p = 0.0068 – one vs. two; p = 0.0774 – one vs. three; p = 0.9335 – two vs. three, ANOVA, Tukey’s post hoc). **(E)** Relationship between the average PSD-95 nanocluster (NC) volume per spine and VGluT2^+^ spine size in apical (R^2^ = 0.1802, slope = 0.0343, 95% confidence: 0.0193-0.049, n = 96 spines) and basal (R^2^ = 0.4235, slope = 0.0658, 95% confidence: 0.0469-0.0846, n = 68 spines) dendrites (**p = 0.009, ANCOVA). **(F)** Average total PSD-95 volume in apical and basal VGluT2^+^ spines with one (L1-3: n = 49 spines, L5: n = 44 spines, p = 0.5569, t-test), two (L1-3: n = 39 spines, L5: n = 21 spines, **p = 0.0047, t-test) and three (L1-3: n = 8 spines, L5: n = 3 spines, *p=0.0476, Student’s t-test) PSD-95 nanoclusters. The total PSD-95 volume per spine was calculated by summing volumes of all PSD-95 nanoclusters (NC) in a single spine. **(G)** Model of trans-synaptic PSD-95/Bassoon nano-organization in VGluT1^+^ and VGluT2^+^ spines in basal and apical compartments of L5 neurons. Bars represent the mean ± SEM acquired from spines on 37 apical and 19 basal dendritic segments of L5 neurons from two biological replicates. Dots on graph indicate individual PSD-95 nanoclusters.

To examine this point in more detail, we correlated average sizes of PSD-95 nanoclusters per spine as a function of increasing spine volume, regardless of spine nanocluster number (Fig. 8e). We reasoned that the low correlation between spine size and nanocluster size would be indicative of more unitary nanoclusters that would be consistent with modular nano-organization. In contrast, a high positive correlation would suggest that changes in nanocluster sizes and spine volumes are linked, indicative of non-modular organization. In line with this idea, we observed significantly higher correlation between average sizes of PSD-95 nanoclusters and spine volumes in L5A VGluT2^+^ spines (R^2^ = 0.4235, slope = 0.065, 95% confidence 0.044 to 0.084) compared to L1-3 VGluT2^+^ spines (R^2^ = 0.1802, slope = 0.034, 95% confidence 0.019 to 0.049, p = 0.009, ANCOVA, Fig. 8e). Despite this observation, there was still a positive correlation between sizes of PSD-95 nanoclusters and VGluT2^+^ spines in L1-3, indicating that PSD-95 nanoclusters in these spines also increase in size as a function of increasing spine volume. However, sizes of PSD-95 nanoclusters in apical VGluT2^+^ spines exhibit a constrained upper limit, suggesting a regulation of their growth across spine sizes and nanomodule numbers. Such restricted growth of individual PSD-95 nanoclusters resembles the modular nano-organization of VGluT1^+^ synapses [18, 19].

Sizes of spines are tightly linked to sizes of PSDs, where large spines with large PSDs are thought to correlate with strong synapses [60, 61]. We therefore asked how total post-synaptic sizes of VGluT2^+^ spines in L5A and L1-3 change as a function of their PSD-95 nanocluster number. Total post-synaptic sizes were calculated by summing volumes of all PSD-95 nanoclusters per spine (Fig. 8f). Single PSD-95 nanocluster-containing VGluT2^+^ spines from both L5A and L1-3 exhibited similar total PSD-95 volumes, which in these spines were equivalent to sizes of single PSD-95 nanoclusters (Fig. 8f, p = 0.5191, t-test). As expected, the total PSD-95 volumes of VGluT2^+^ spines with two PSD-95 were larger than those with one PSD-95, and VGluT2^+^ spines with three PSD-95 had larger PSD-95 volumes than spines with two PSD-95 nanoclusters in both layers (Fig. 8f, p < 0.0001, ANOVA). The increase in total PSD-95 sizes as a function of PSD-95 nanocluster number was more robust for VGluT2^+^ spines in L5A. Consistent with the model that sizes of individual PSD-95 nanoclusters scale with increasing volumes of VGluT2^+^ spines on basal but not apical dendrites, two PSD-95 nanocluster-containing VGluT2^+^ spines from L5A had significantly larger total PSD-95 sizes than L1-3 VGluT2^+^ spines with equivalent numbers of PSD-95 nanoclusters (Fig. 8f, p = 0.0017, t-test). Similarly, L5A VGluT2^+^ spines with three PSD-95 nanoclusters had significantly larger total PSD-95 sizes than their VGluT2^+^ spine equivalents in L1-3 (Fig. 8f, p = 0.0476, t-test). These data indicate that VGluT2^+^ spines on basal dendrites assemble post-synaptic macrostructures from larger nanoscale building blocks and thus exhibit structural correlates of stronger synapses than their counterparts in apical dendritic compartments [60, 61].

## Discussion

A fundamental challenge of modern neuroscience for understanding brain function is deciphering how rich molecular and structural diversity of active zones and PSDs endow synapses with specific functions that underlie information storage and processing in neural circuits. How functionally diverse synapses organize their nano-architecture is poorly understood due to difficulties in obtaining three-dimensional reconstructions of individual synapses in complex brain tissue with the resolution required to visualize the organization of pre- and post-synaptic molecular complexes. Combining nanobodies with five-channel confocal and 3D-STED imaging of individual TC and CC synapses on L5 pyramidal neurons in S1, we show that there is a degree of variability in how functionally diverse glutamatergic synapses in the brain are organized at the nanoscale. Compared to CC synapses that predominantly contained only one trans-synaptic nanomodule (66%), TC afferents in apical compartments formed on spines with single (51%) and multiple (47%) aligned PSD-95 and Bassoon nanoclusters with a similar frequency. However, the sizes of individual PSD-95 nanoclusters remained relatively stable between single and multi-clustered spines, indicating that apical TC spines follow modular principles of nano-organization. In TC synapses on basal dendrites, sizes of PSD-95 nanoclusters scaled with the increasing size of VGluT2^+^ spines, deviating from modularity, and only very large spines were multi-clustered. These results shed light on the importance of the type (VGluT1 vs. VGluT2) and location (apical vs. basal) of afferent input in shaping synaptic nano-architecture in the brain (Fig. 8g).

Super-resolution imaging in intact brain tissue is challenging. Resolution declines with tissue depth, limiting super-resolution only to <10 µm from the surface [44]. STED nanoscopy is well suited for probing nanoscale organization of synapses in the brain due to its near-infrared (775 nm) STED that attenuates light absorption and scattering. STED allows fast simultaneous imaging of multiple fluorophores without compromising resolution between channels with minimal chromatic aberration in XY and Z planes [18]. Using tau-STED, which combines optical signals from conventional STED with physical information of fluorescent lifetimes of individual photons [53], we were able to resolve objects with a lateral separation of 50 nm and axial separation of 90 nm without the need for deconvolution. Together with a recent study applying STED nanoscopy to obtain molecularly informed synapse-level volume reconstruction in living brain tissue, these data suggest that <100 nm X, Y, Z resolution needed for 3D reconstruction of synapses and their molecular constituents in the brain is attainable using the tau-STED approach that mitigates photobleaching and improves resolution [62]. Importantly, our STED imaging of fixed brain cryosections enables simultaneous visualization of multiple endogenous molecules to assign their nano-organization to spines of varying morphologies.

Precisely aligned nanodomains consisting of proteins in active zones and PSDs form trans-synaptic units that are key features regulating synapse function [15, 18, 19, 22]. Thus, identifying both pre- and post-synaptic molecules is critical for the accurate reconstruction of synaptic nano-architecture in the brain’s dense neuropil. However, synaptic specializations are poorly accessible to conventional antibodies in paraformaldehyde fixed samples [43, 49]. Indeed, the PSD-95 antibody fails to identify “real” post-synaptic clusters from random cluster simulations without subjecting the tissue to antigen retrieval. In contrast, the same tissue treatment negatively affects pre-synaptic VGluT1 and Bassoon antibody labeling. As a result, colocalization between PSD-95 and VGluT1 or Bassoon clusters visualized by antibodies identified only ∼17% to 25% of putative excitatory synapses and 30% of trans-synaptic nanomodules in brain sections. Nanobodies significantly improved the labeling of synaptic molecules in paraformaldehyde fixed brain sections without a need for tissue modification. The effect of nanobodies on PSD-95 labeling that localizes to PSDs was most apparent, while the labeling for VGluT1 was comparable between nanobodies and antibodies. Critically, the density of colocalized nanobody-labeled PSD-95 and antibody-labeled Bassoon much closer approximated dendritic spine densities in dendrites of Thy1-YFP-H mice than the density of colocalized puncta visualized by staining purely with conventional antibodies. However, the most essential feature of nanobodies is their ability to visualize diffraction-limited synaptic clusters *in vitro* and in brain sections.

Despite these advantages, the use of nanobodies also presents certain limitations. Since nanobodies are not subjected to the secondary antibody signal amplification, clusters exhibit low brightness. We have chosen a highly abundant synaptic molecule – PSD-95 to characterize the nanobody efficiency for STED nanoscopy *in situ*. Visualizing AMPARs and NMDARs that are less abundant at PSDs could suffer from low signals that may compromise their ability to be precisely localized *in vivo*. We could not address the efficacy of nanobodies targeting less abundant synaptic molecules for STED imaging due to a limited repertoire of available nanobodies targeting endogenous synaptic proteins. The emergence of sequences for recombinant monoclonal antibodies should accelerate the purification and screening of nanobody-like single-chain variable domains (scFv) for labeling receptors and other synaptic molecules for super-resolution imaging in the brain [45, 63]. Additionally, orthogonal labeling using non-canonical amino acids that sidestep antigen masking inherent to proteins in PSDs has been recently shown to be a useful tool for super-resolution imaging [64]. Alternatively, CRISPR gene editing that enables direct fusion of fluorescent probes to endogenous proteins could facilitate super-resolution imaging of synaptic molecules in the brain, although such approaches suffer from their own limitations [65, 66]. Finally, non-aldehyde fixation protocols should improve antigen accessibility, further enhancing the labeling efficiency of synaptic proteins in the brain [14].

Much of our understanding of the molecular nano-organization of the synapse comes from the exploration of synaptic nanoarchitecture of neurons in cell culture. Super-resolution imaging of glutamatergic and GABAergic synapses revealed that trans-synaptic nano-organization follows conserved principles, with an ordered network of aligned pre- and post-synaptic proteins regulating synaptic transmission and plasticity [17–23, 25]. At excitatory synapses, post-synaptic PSD-95, AMPARs, and NMDARs and pre-synaptic Bassoon, VGluT1, and SYT1 share a modular organization, where the number but not the size of precisely aligned pre- and post-synaptic nanoclusters increases with spine size [18, 19]. Previous work showed that the nanoscale organization of AMPARs and NMDARs containing specific subunits differs across spines with a different number of nanomodules resulting in synapses with non-overlapping AMPAR and NMDAR nanodomains [17, 18, 67]. Our findings demonstrating that apical and basal TC synapses exhibit variable nano-architecture with different numbers and sizes of PSD-95 nanoclusters raise a question of how AMPAR and NMDAR nanodomains are organized in these synapses. Additionally, SAP-102 and PSD-95 MAGUK family members that differentially regulate AMPAR and NMDAR trafficking to synapses exhibit distinct expression in VGluT1^+^ and VGluT2^+^ spines [13, 68]. Thus, it remains to be determined whether AMPAR and NMDAR sub-types exhibit different nano-organization in TC vs. CC synapses and how it might impact the function of these two diverse glutamatergic synapses.

It has long been proposed that TC inputs to L4-6 function as glutamatergic drivers, whereas TC inputs to L2-3 are glutamatergic modulators [69, 70]. TC synapses on basal dendrites have larger PSD-95 nanoclusters than TC synapses in L1-3.

Furthermore, VGluT2^+^ spines on basal dendrites with two and three PSD-95 nanoclusters have significantly larger total PSD-95 volumes than VGluT2^+^ spines with the equivalent number of PDS-95 nanoclusters in apical compartments. The large post- synaptic nanoscale features of L5 TC synapses are consistent with work showing relatively strong TC synaptic currents on basal dendrites of L5 neurons [55]. Selective enlargement of L5A inputs and associated PSD-95 has been recently linked to experience-dependent plasticity [71]. It will be essential to determine whether TC synapses on basal dendrites lack mGluR nanodomains, a characteristic feature of glutamatergic drivers [70]. In contrast, many VGluT2^+^ spines in L1-3 contain multiple aligned PSD-95/Bassoon nanomodules. The presence of multiple pre-synaptic nanoclusters on a single TC spine implicates a mechanism that could impact the quantal mode of neurotransmitter release [4, 36]. Previous work in cell culture demonstrated that the number of aligned nanomodules rapidly increases after chemical LTP [19]. An ability to dynamically adjust synaptic nanomodule numbers could provide a means for rapid tuning of synaptic transmission needed to modulate neuronal activity.

Indeed, PSD-95 sizes in L1 are rapidly reduced, likely as a way of compensating for the enlargement of PSD-95 areas on spines in L5A during learning [71]. The structure-function relationship of TC synapses remains enigmatic. Despite their lower abundance compared to CC synapses, TC synapses are better at activating cortical neurons. The reason for this is difficult to unlock, given that very few studies directly compare the function of TC and CC synapses, leading to controversial theories. Early *ex vivo* studies suggested that TC synapses release multiple quanta and are more reliable than CC synapses [36]. Others have shown that TC synapses are stronger than CC synapses [37, 38, 58]. Alternatively, convergence of TC inputs that synchronizes cortical firing has been proposed [39–41]. Recent work examining L2/3 neurons in the visual cortex suggested that TC inputs favor smaller spines than CC inputs, which provides high readout accuracy within the heterogenous network of L2/3 neurons [42].

Using a vast dendritic tree of genetically identified L5 neurons as a model, our super-resolution approach reveals laminar differences in how TC synapses are organized. Remarkably, CC synapses in S1 do not exhibit nanoscale laminar variation. Therefore, it might be challenging to establish a unifying principle that could generalize the contribution of diverse TC synapses to processing of sensory information, even in the same neuron. The idea of laminar diversity of glutamatergic synaptic nanoarchitecture is consistent with the recent work demonstrating that the nanoscopic landscape of critical regulators of synaptic specification that bind Neurexins, Neuroligins and LRRTMs, differs across TC and CC inputs and brain regions [72]. Future studies are needed to determine how input/target selectivity might engage different trans-synaptic and cell-autonomous mechanisms to establish diverse synaptic nanodomains that may create unique signaling environments for synaptic processing and cortical computation in various sub-cellular compartments and cortical layers.

## Materials and Methods

### Animals

All animal studies were approved by the Institutional Animal Care and Use Committee guidelines at West Virginia University (#2102040142_R1 and #2105042129) in accordance with the US National Institutes of Health guidelines. Thy1-YFP-H mice (B6.Cg-Tg(Thy1-YFP)16Jrs/J, Strain # 003782, RRID: IMSR_JAX:003782) were obtained from Jackson Laboratory and housed (2-5 mice per cage) in West Virginia University Health Sciences Center’s laboratory animal facility. Both male and female mice were used for experiments at 1-3 months of age. Timed pregnant Long-Evans rat females used for the collection of E17-18 male and female rat embryos to make primary cortical neuron cultures were purchased from Charles River Laboratories Inc. (Wilmington, MA).

### Brain tissue collection and processing

To procure mouse brain sections, mice were perfused using previously described methods [19]. Briefly, mice were anesthetized with 4% isoflurane until unable to right themselves and first perfused transcardially at a rate of 7mL/minute for 2 minutes with ice-cold ACSF (125 mM NaCl, 2.5 mM KCl, 2 mM CaCl2, 25 mM NaHCO3, 1.25 mM NaH2PO4, 2 mM MgSO4, 10 mM Glucose, and 0.052 mg/mL Heparin, pH 7.4), followed by 60mL ice-cold 4% paraformaldehyde (PFA) (ref # 00380-1, Polysciences, Warrington, PA) in 0.1 M phosphate buffer (80 mM Na2HPO4 and 19 mM NaH2PO4, pH 7.4) containing 0.003% glutaraldehyde (ref # 1875, VWR, Radnor, PA). Brains were removed and post-fixed overnight at 4°C in 4% PFA. After washing 3 times (10 minutes each wash) in 1xPBS (ref # 70011-044, Gibco, Jenks, OK), brains were transferred to 15% sucrose solution and allowed to equilibrate overnight at 4°C, then transferred to 30% sucrose solution and allowed to equilibrate at 4°C for up to 48 hours. In preparation for cryosectioning, brains were embedded in Tissue-Tek O.C.T. Compound (ref # 4583, Sakura Finetek) by submerging it in 2-Methylbutane (ref # O3551-4, Fisher Scientific, Pittsburgh, PA) cooled with dry ice to −80°C. Brains were sectioned at 50 µm or 6 μm on a Leica CM3050 S cryostat set to −20°C for both the object and chamber temperature (Leica Biosystems, Nussloch, Germany). We collected sections from the primary somatosensory cortex (coronal sections from bregma: A/P −1.05 mm to −2.1 mm). Free-floating 50 μm sections were kept in 1xPBS plus 0.01% sodium azide until use. Thin 6 μm sections were mounted on pre-treated (5 mg/mL gelatin from bovine skin and 1 mM chromium (III) potassium sulfate) 1.5 coverslips and used immediately for immunohistochemistry.

### Immunohistochemistry of brain sections

Brain sections underwent one of the following immunohistochemistry (IHC) protocols:

#### No treatment

Brain sections were blocked and permeabilized for 1 hour in 1xPBS containing 5% normal goat serum (ref # 16210064, Thermo Fisher Scientific) and 0.5% Triton X-100 (ref # BP151, Thermo Fisher Scientific). Tissue was incubated in primary antibodies or nanobodies (see Table 1 for details) diluted in blocking buffer for 48 hours at 4°C. Tissue was washed three times with 1xPBS (10 min each wash) and then incubated in secondary antibodies (Table 1) for 2 hours at room temperature. After washing 3 times with 1xPBS, brain sections were mounted in ProLong Glass Antifade mounting medium (ref # P36984, Invitrogen, Waltham MA) and kept at room temperature until imaging.

**Table 1.**
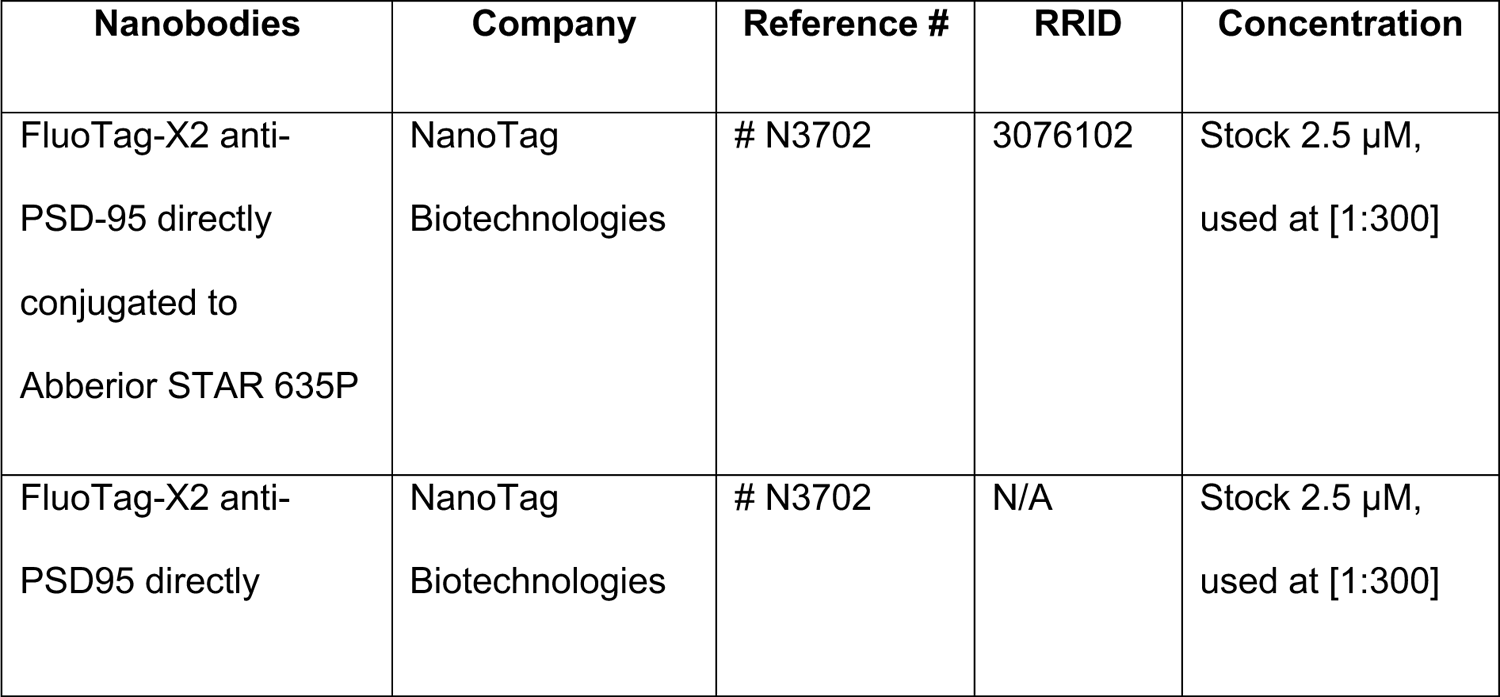
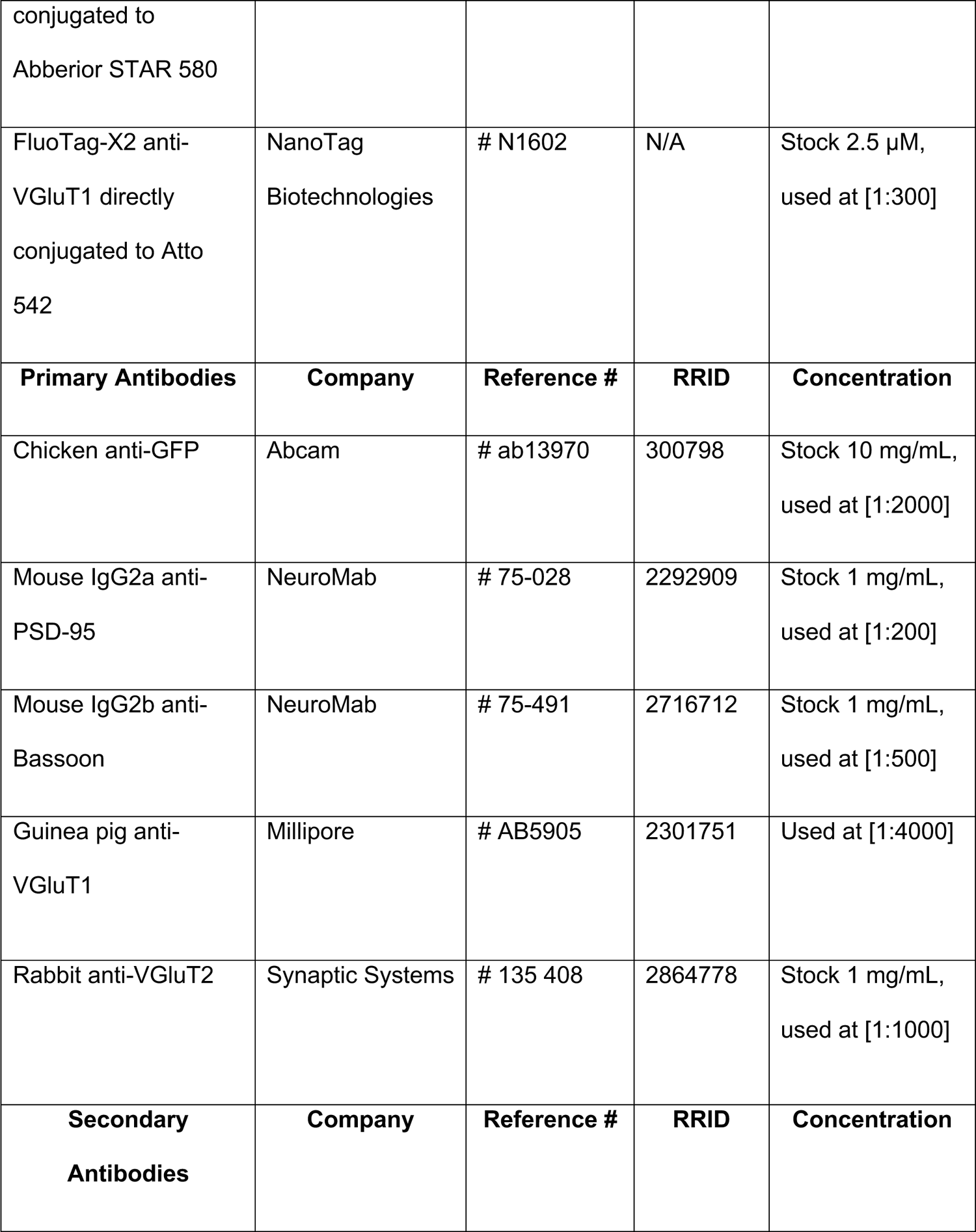
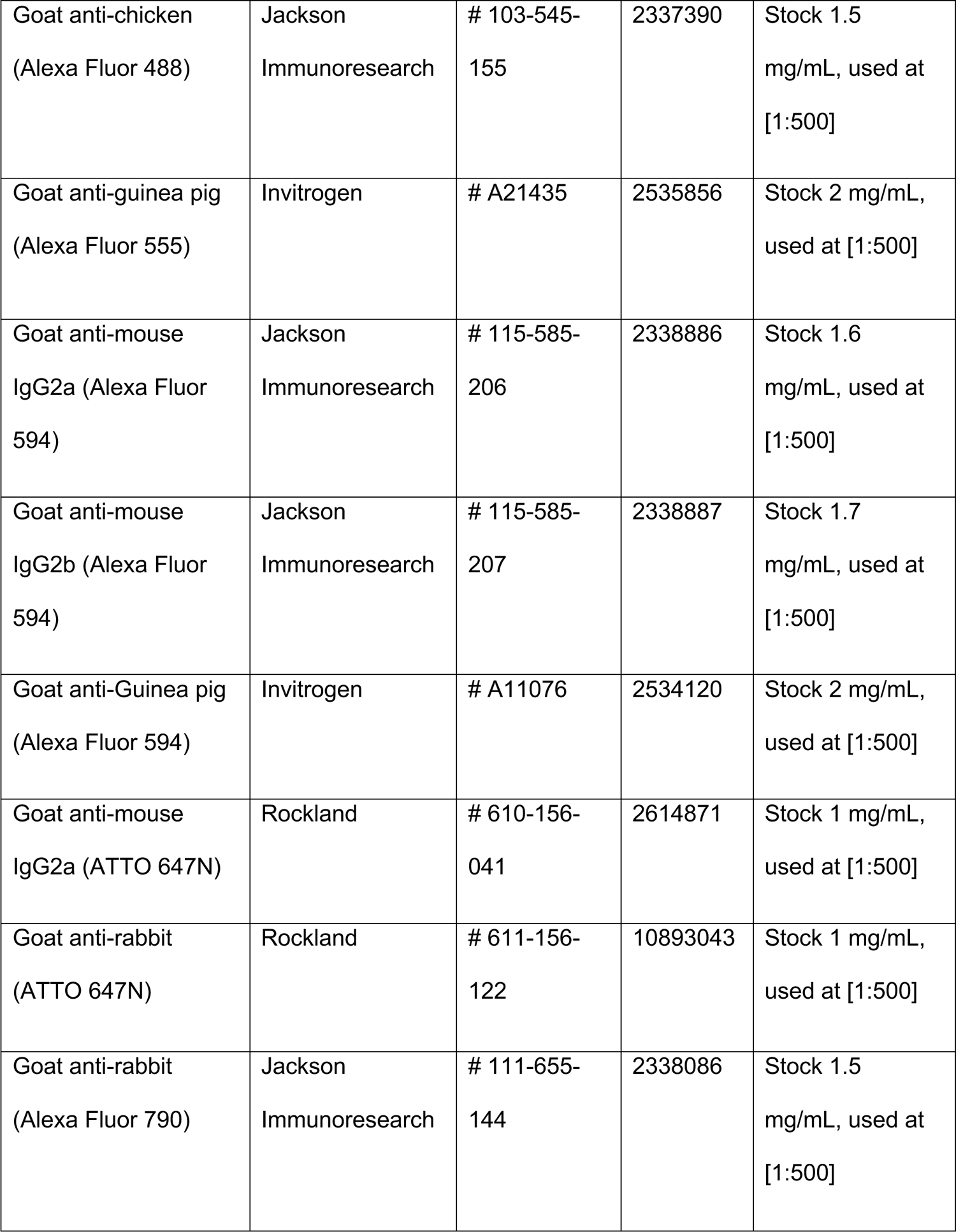
Nanobodies and Antibodies.

#### Antigen retrieval

Before the IHC, brain sections were first subjected to a 5 minute incubation in MiliQ water with gentle rocking at 37°C, followed by either a 2 minute (6 µm sections) or 10 minute (50 µm sections) rocking at 37°C in pepsin reagent (ref # R2283, Millipore Sigma, Burlington, MA). The pepsin reaction was quenched by successive incubations in blocking buffer. The tissue was then subjected to immunostaining with primary and secondary antibodies as described above for the no treatment protocol.

#### CUBIC (Clear Unobstructed Brain Imaging Cocktails)

CUBIC clearing of brain sections was performed as previously described [19, 51]. Briefly, 50 µm and 6 µm brain sections were blocked overnight in blocking buffer, followed by a 4 day incubation at 4°C in primary antibodies diluted in blocking buffer. Sections were washed with 1xPBS (3 times for 10 minutes) and then incubated overnight in secondary antibodies diluted in blocking buffer. After washing 3x in 0.1M phosphate buffer (PB), tissue was placed in the CUBIC reagent, consisting of 25% w/v urea (ref # U5378, Millipore-Sigma, St. Louis, MO), 25% w/v Quadrol (ref # 122262, Millipore-Sigma), and 0.2% w/v Triton X-100 (ref # T8787, Millipore-Sigma). Upon visible clearing of tissue (30 minutes for 50 µm sections and instantly for 6 µm sections), tissue was mounted in the CUBIC reagent, slides were sealed with nail polish and imaged within 48 hours.

#### CUBIC + antigen retrieval

The tissue first underwent the antigen retrieval protocol using pepsin reagent as described above, followed by immunostaining and tissue clearing described for the CUBIC protocol. Sections were mounted in the CUBIC reagent, and slides were sealed with nail polish and imaged within 48 hours.

### Primary cortical neuron culture preparation

Dissociated cortical neurons were prepared from the embryonic day 17-18 (E17-18) male and female rat cerebral cortexes as described previously [18, 19]. Primary cortical neurons were cultured for 21-25 Days In Vitro (DIV) in Neurobasal medium with phenol red (cat#: 12348017, Thermo Fisher Scientific) supplemented with 1x B27 supplement (ref # 17504044, Thermo Fisher Scientific), 200 mM L-glutamine (cat#: 25030081, Thermo Fisher Scientific) and 0.1 mg/mL penicillin-streptomycin (ref # 15140122, Thermo Fisher Scientific). Neurons were plated on poly-D-lysine (ref # 354210, Corning, Corning, NY) and laminin (mouse, ref # 354232, Corning) coated glass coverslips (12 mm, #1.5; ref # 64-0732, Warner Instruments, Camden, CT). Neurons were plated at 150,000 cells/well in 24-well plates and were maintained in a humidified 37°C incubator with 5% CO2 until they were used for imaging at DIV 21-25.

### Neuronal transfection

Neurons were transfected at DIV 3 as previously described [18, 19] using Lipofectamine 2000 (cat#: 11668027, Thermo Fisher Scientific). EGFP, under the control of a human ubiquitin promoter (pFUg-EGFP), was used as a cell filling dye to visualize neuronal morphology [18, 19]. Briefly, for each well of a 24-well plate, the conditioned medium was first collected from plated neurons and replaced with 300 µl of Neurobasal medium without any supplements. 100 µl of transfection mix containing 0.5 µl of Lipofectamine 2000 and 200 ng of pFUg-EGFP plasmid was then added to each well of a 24-well plate. Neurons were incubated with the transfection cocktail at 37°C for 2 hours. After 2 hours, the transfection medium was replaced with 500 µl of the warmed conditioned medium that was sterilized by passing through a 0.2 µm filter (Millipore-Sigma). Transfected neurons were then placed in a humidified 37°C incubator until DIV 21-25, at which point they were used for immunocytochemistry and STED imaging.

### Immunocytochemistry

Cultured cortical neurons were fixed between DIV 21 and DIV 25 in 4% PFA containing 2% sucrose supplemented with 0.003% (v/v) glutaraldehyde in 1xPBS for 8 minutes at room temperature [18, 19]. Fixed neurons were then washed three times in 1xPBS. Coverslips were next blocked and permeabilized for 1 hour at room temperature in 1% ovalbumin (ref # A5503, Millipore-Sigma) and 0.2% gelatin from cold-water fish (ref # G7041, Millipore-Sigma) in 1xPBS containing 0.1% saponin (ref # 558255, Millipore-Sigma). Neurons were immunostained for 2 hours at room temperature or overnight at 4°C with the indicated primary antibodies or nanobodies, washed three times in 1xPBS, and then exposed to corresponding secondary antibodies (if needed) for 1 hour at room temperature. After washing three times in 1xPBS, coverslips were mounted with ProLong Glass antifade mounting medium and used for confocal and STED imaging after 24-48 hours, when the mounting medium was fully cured.

### Confocal imaging and analysis

50 µm and 6 µm sections collected from Thy-1-YFP-H mouse somatosensory cortex (S1) were imaged by confocal scanning microscopy using a Leica Stellaris 8 confocal and STED microscope system (Leica Microsystem, Mannheim, Germany). Confocal images were acquired with a 100x oil immersion objective (1.4 Numerical Aperture (NA), Leica) at 1.7-1.9 zoom to obtain a 100 nm pixel size. Stacks were the collection of 10-12 images taken at 0.3 µm intervals and were analyzed using ImageJ as maximum intensity projections with the experimenter blinded to treatment conditions. For each experimental condition, images were taken from at least three different Thy-1-YFP-H animals.

### Puncta co-clustering analysis

Analyses were performed blinded to the experimental condition (treatment of brain tissue). Individual pre- and post-synaptic puncta and pre- and post-synaptic co-clusters were analyzed using ImageJ custom-build macros designed to automatically count the number and size of clusters along GFP labeled dendrites as described previously [73–76]. Each 512 x 512-pixel channel was thresholded to the mean + 2 x standard deviation (SD) and converted to a binary image. Individual pre- and post-synaptic puncta were defined as continuous clusters of 3-100 pixels (the size of each pixel was 100 nm) along at least 50 µm of dendrite. For the colocalization analysis, co-clusters were defined as > 1-pixel overlap between binarized pre- and post-synaptic clusters from two different channels.

### Monte Carlo analysis

Using PSD-95, VGluT1 and Bassoon cluster numbers and sizes obtained from the unbiased measurements of cluster densities (described above), we simulated random cluster distributions within the 512 x 512-pixel frame using a random cluster generator plug-in in ImageJ [19]. We then modeled probabilities of overlap between randomly generated clusters and YFP-labeled dendrites and dendritic spines by overlaying simulated clusters onto raw YFP images obtained from confocal imaging. Individual cluster sizes were approximated by a circular area calculated from the average PSD-95, VGluT1 and Bassoon cluster areas obtained from our confocal images. Densities of randomly generated PSD-95, VGluT1 and Bassoon clusters and co-clusters along the YFP labeled dendrites were determined in an unbiased manner using ImageJ custom-built macros described above [73–76]. Simulations were conducted for every YFP-labeled dendritic segment collected from confocal imaging.

### STED imaging

Two-color tau-STED imaging of endogenous proteins was performed on fixed neurons in cell culture and 6 µm brain sections collected from Thy-1-YFP mice based on protocols developed previously [18, 19]. A Leica Stellaris 8 3D tau-STED confocal and super-resolution system (Leica Microsystem, Mannheim, Germany) equipped with a tunable white light laser (WLL), CW 592 nm, CW 660 nm and pulsed 775 nm STED depletion lines was used for image acquisition. HyD-X detectors set to photon counting mode at a 12 mV threshold were used to capture single photons for STED image acquisition. The 100x oil immersion objective (1.4 NA) with 4.5x digital zoom to obtain ∼20 nm pixel size was used to acquire image stacks at 150 nm intervals using 400 Hz scanning of 1024 x 1024-pixel imaging fields (∼ 20 µm x 20 µm). Line accumulations (2x – 3x) were used to image individual channels. Stacks for each channel were acquired in a sequential mode using the “between stacks acquisition” setting. Target proteins labeled with Abberior STAR 635P, Atto 647N, Alexa Fluor 594, Abberior STAR 580, dyes directly conjugated to nanobodies, or secondary antibodies (Table 1) were acquired using the Fast Lifetime Contrast (FALCON) enabled tau-STED module (Leica) with the time-gate on HyD-X detectors adjusted between 0.1 and 6 nanoseconds.

Abberior STAR 635P/Atto 647N labeled endogenous proteins were excited with the 635 nm laser (10-15% maximal AOBS laser power). Alexa Fluor 594/Abberior STAR 580 labeled endogenous proteins were excited using the 594 nm laser (5-12% maximal AOBS power). The pulsed 775 nm STED depletion line set at 10% of maximal AOBS laser power was used for STAR 635P/Atto 647N fluorophores and 15% of maximal AOBS laser power for Alexa Fluor 594/Abberior STAR 580 fluorophores to generate STED. Using these settings, we were able to obtain an XY resolution of ∼50 nm, measured by FWHM of GATTA STED nanorulers (STED 50R Brightline, GATTAquant, GMBH, Munich, Germany, Fig. S2) [18]. All data shown were imaged using 3D STED with 20% STED laser power re-directed toward the Z-doughnut, which allowed us to resolve GATTA 3D STED nanorulers spaced 90 nm along the Z-axis (STED 3D 90R, GATTAquant, Fig. S2). For simultaneous confocal/STED imaging, fluorophores that were not exposed to STED lasers (EYFP, Alexa Fluor 790, Alexa Fluor 555) were imaged first, using only excitation lines provided by the WLL. This was then followed by STED acquisition of Abberior STAR 635P/Atto 647N fluorophores and then by STED acquisition of Alexa Fluor 594/Abberior STAR 580 fluorophores, all in the same sequence.

### Image processing

Following image acquisition, background photons were removed from raw STED images by adjusting tau strength using the tau-STED module in the FALCON application on the Stellaris 8 STED instrument (Leica Microsystems). The ability to remove unwanted photons relies on generating fluorescence lifetime imaging (FLIM) profiles of every photon captured by HyD-X detectors [53]. The tau strength was adjusted to reduce blur surrounding cluster edges in a manner that did not distort existing clusters or create cluster artifacts. With 5-10 captured photons per pixel, this usually resulted in a tau strength between 100-200. Once the tau strength was determined for each STED channel, it was kept constant throughout all experiments.

Afterward, tau-STED images were subjected to 0.8-pixel Gaussian blur, and brightness was adjusted to generate high-contrast images for figures and analysis. None of the images acquired were deconvolved. Tau-STED images were obtained as 16-bit Tiff files. These images were initially adjusted in ImageJ by subtracting the background (mean pixel intensities of the entire 1024 x 1024 frame) from every channel. We then converted these 16-bit files to 8-bit images that were subsequently used for 3D reconstructions in the Neurolucida 360 software (MBF Bioscience, Williston, VT).

### STED analysis

All analyses were conducted offline using Fiji ImageJ and built-in macros, as described below. Neurolucida 360 (MBF Bioscience) was used to reconstruct confocal and STED images in 3D and obtain parameters pertaining to individual clusters and spines.

#### Nanomodule identification

Super-resolution analysis of synaptic cluster localizations in dendritic spines was performed on a per spine basis as described [18, 19]. Images of YFP labeled spines, VGluT1 and VGluT2 were acquired at confocal resolution (∼250 nm), and a 0.8-pixel Gaussian blur was applied to filter out noise.

Individual spines were converted to binary masks by thresholding YFP (enhanced by GFP immunolabeling) to the mean +2 x SD of the 1024 x 1024-pixel area corresponding to the entire image field. The localization of PSD-95 and Bassoon nanoclusters was then determined per spine using two different approaches. First, ROIs (regions of interest) of each thresholded spine head were used to manually assign STED-resolved puncta to spines. PSD-95 clusters were assigned to a spine if the thresholded pixel areas were entirely within the spine head ROI. Bassoon clusters were assigned to a spine if the thresholded pixel areas either entirely or partially overlapped with the spine head ROI. Spine ROI colocalization of each cluster was made independently for each Z-section. Orthogonal views of overlaid image stacks were used to verify that individual clusters colocalized with spine ROI in the Z-plane. Following the nanocluster assignment, we categorized spines either as VGluT1 or VGluT2 based on their colocalization (> 1 pixel overlap) with YFP-labeled dendritic spines. Sometimes, both VGluT1 and VGluT2 appeared to colocalize with the same spine. In those cases, we used the colocalization of aligned nanoscale-resolved Bassoon and PSD-95 with either VGluT1 or VGluT2 to determine under which glutamatergic marker to categorize spines with apparent dual innervation.

In our second approach, we used the 3D visualization and analysis software Neurolucida 360 (MBF Bioscience) to independently verify the 3D segmentation of individual nanoclusters by reconstructing confocal and tau-STED channels. We converted 16-bit images to 8-bit Tiff files by assigning the value of 255 to the maximum pixel intensity value in 16-bit images. We then reconstructed dendritic trees by tracing GFP-filled dendrites using the Rayburst crawl mode [77]. Spines were detected automatically by adjusting the outer range to a 2.5 µm maximum value, a minimum height of 0.3 µm, a detector sensitivity of 50-150%, and a minimum count of 10 voxels. These values were sufficient to reconstruct most spines. A minority of spines had to be reconstructed manually by adjusting the above values. PSD-95 and Bassoon nanoclusters were reconstructed in 3D using detector settings in Neurolucida 360 with a diameter of 0.1-0.5 µm and a sensitivity set to 200-300%. Clusters were reconstructed only if they contained a minimum of 10 contiguous voxels. 3D-resolved PSD-95 nanoclusters were then assigned to individual spines in a semi-automatic manner by setting the proximity value between nanoclusters and spine heads to 0.01 µm (inside the spine). For Bassoon, this value ranged from 0.15-0.2 µm. Some clusters had to be reconstructed manually by increasing either detector diameter, sensitivity or both.

Reconstructed images were then exported to the associated 3D Neurolucida Explorer (MBF Bioscience) to obtain the measurements (volumes and nanocluster numbers per spine) for 3D reconstructed clusters.

#### 3D STED cluster distance analysis

Colocalization and nearest neighbor analysis of 3D STED-resolved synaptic clusters were performed on a per-cluster basis using the entire area of an image (1024 x 1024-pixel format). Segmentation and subsequent measurements of distances of segmented clusters were performed using the DiAna plug-in in ImageJ that enabled the analysis to be done in an automated way [54]. 3D-resolved nanoclusters of synaptic proteins (acquired in tau-STED super-resolution) were identified by binarizing each channel separately using intensity thresholds followed by segmentation of individual clusters in the DiAna segmentation window [18].

Segmentation of all synaptic clusters was performed using the local maxima method combined with user-defined thresholds [18, 54]. Local maxima were identified using a radius of 5 pixels (100 nm) in the XY-plane and 2 pixels (300 nm) in the Z-plane. Local thresholds were defined by thresholding individual 16-bit images to the mean + 2 x SD of intensity values in an entire 1024 x 1024 image frame. The maximum radius of segmented clusters was set to 20 pixels (individual pixel sizes in our images were set to 20-25 nm to allow a maximum resolution of ∼50 nm). The standard deviation for Gaussian fit and threshold calculation was set to 1.5. Minimum and maximum voxel sizes were set to 3 and 20,000, respectively. Distance analysis of segmented clusters was based on classical Euclidean distance computation [54]. For measurements of the nearest neighbors and trans-synaptic clusters, we implemented center-to-center distances where, for each object from one image (channel 1), the center-to-center distances with all objects from another image (channel 2) were computed in 3D, and closest neighbor distances were reported [18].

### Experimental Design and Statistical analyses

Data were acquired and analyzed based on the standards in the field; however, no method of biological specimen randomization was used to determine how samples were allocated to experimental groups and processed. Unless otherwise stated, data are expressed as means ± SEM. All data points collected were included for analysis. Statistical significance of the differences among groups was determined by a one-way analysis of variance (ANOVA) followed by post-hoc tests described in individual figure legends. When testing differences between two conditions, we performed a two-tailed Student’s t-test. Kolmogorov-Smirnov (K-S) tests were used to test differences between non-parametric probability distributions. P values less than 0.05 were considered statistically significant. For p values less than 0.0001, we provide a range and not the exact number (e.g., p<0.0001). The data distribution was assumed to be normal, but this was not formally tested. Sample sizes were determined based on previous publications [18, 19, 73–76]. All statistical analyses were performed using Graph Pad Prism statistical software (https://www.graphpad.com). Unless stated otherwise, statistical tests were conducted on a per spine basis. Statistical tests for center-to-center distances between 3D STED clusters were performed on a per cluster basis. Data were collected from a minimum of three independent transfection experiments or at least two different adult mice between ages P35-P90.

#### Biological reagents

Following nanobodies, primary and secondary antibodies were used to immunostain brain sections and cultured neurons. Dilution and specificity of all antibodies used were profiled in prior publications and were reported to be specific [18, 19, 73–76]. PSD-95 nanobody has been shown not to cross-react with mouse and rat PSD-93, SAP-97 and SAP-102 [52]. VGluT1 specificity was tested in the current study by determining colocalization with antibodies for VGluT1 and VGluT2 (Figs. 3 and S1).

## Data availability

Data supporting the findings of this study are available within the article, its supplementary information. All data are fully available without restriction.

## Acknowledgments

We thank Dr. Ariel Agmon in the WVU Department of Neuroscience and the members of the Hruska laboratory for their critical reading of our manuscript and helpful suggestions and comments. Grants from NIGMS (P20GM109098) and Alzheimer’s Association (AARG-NTF-23-1150820) to M.H. supported this work. The content is solely the responsibility of the authors and does not necessarily represent the official views of the NIH or Alzheimer’s Association.

## Supplementary Information

**Figure S1.**
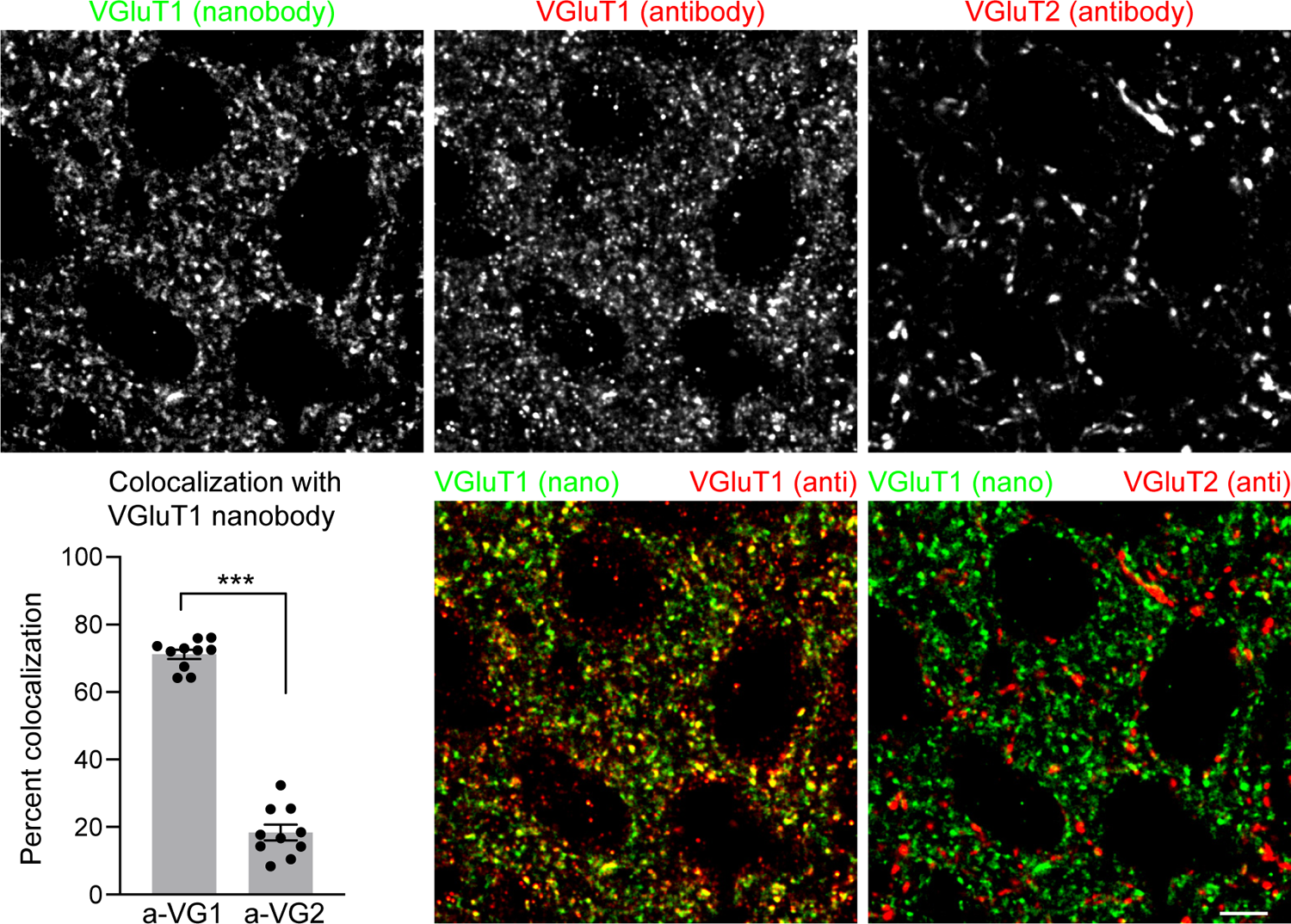
VGluT1 nanobody is specific for VGluT1 but not for VGluT2. Representative confocal images of nanobody-labeled VGluT1^+^ (Atto 542, green), antibody-labeled VGluT1^+^ (Alexa Fluor 594, red) and antibody-labeled VGluT2^+^ (Atto 647N, red) clusters in S1. The graph indicates average percent colocalization between nanobody-labeled VGluT1^+^ clusters and antibody-labeled VGluT1^+^ (a-VG1) or VGluT2^+^ (a-VG2) puncta (n = 10 images, ***p<0.0001, t-test). Bar graphs represent mean ± SEM. Dots on bar graphs represent data from individual image fields collected from at least 3 independently immunostained brain sections. Scale bar, 5 µm.

**Figure S2.**
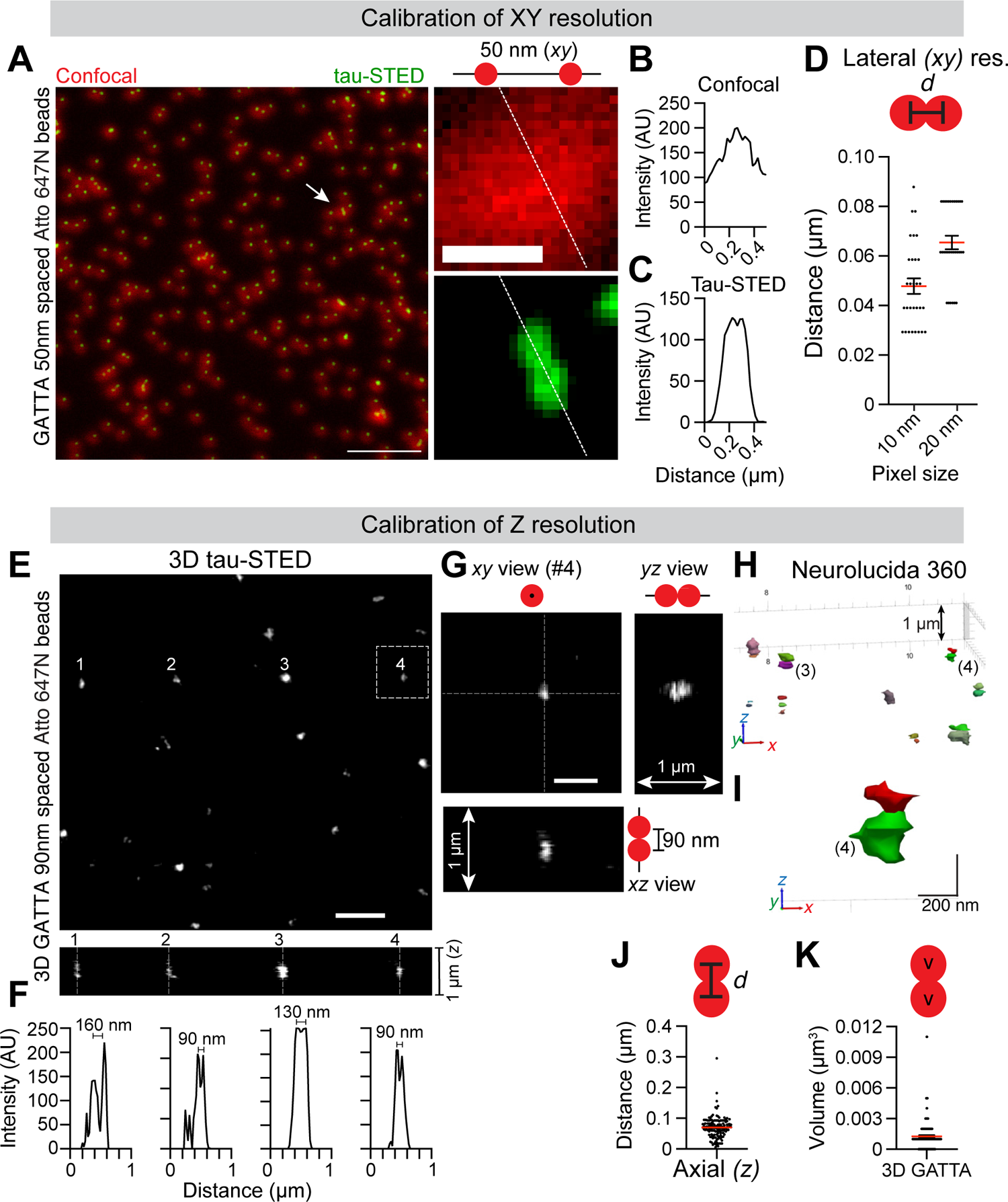
Calibration of 3D tau-STED resolution. **(A)** Representative image of GATTA 50 ± 5 nm laterally spaced Atto 647N-labeled nano ruler beads imaged in confocal (red) and tau-STED (green). Arrow indicates a single GATTA bead, shown in a larger view in confocal and STED channels on the right. Scale bars, 2 µm (left), 250 nm (inset, right). **(B, C)** Line profiles through a single GATTA bead from the inset in **a** (white dashed line) imaged in the confocal (**B**) and tau-STED (**C**) resolution. **(D)** Average peak-to-peak distance calculated from line profiles of 50 ± 5 nm spaced GATTA beads. Measurements were made for image formats with the 10 nm (d = 47.8 ± 3 nm, n = 30 beads) and 20 nm pixel size (d = 65.5 ± 2.7 nm, n = 31 beads). **(E)** Representative image of 3D GATTA 90 ± 5 nm axially spaced Atto 647N-labeled nano ruler beads imaged in 3D tau-STED. The bottom image shows orthogonal *xz* views of beads # 1-4. Scale bar, 1 µm. **(F)** Line profiles of beads # 1-4 in the *xz* view (shown in **e**) demonstrating peak-to-peak separation of 90 nm to 160 nm. **(G)** Zoomed-in view of the bead # 4 and accompanying orthogonal (*xy* and *xz*) projections. Scale bar, 500 nm. **(H, I)** 3D Neurolucida 360 reconstruction of GATTA beads from **E**. Beads #3 and #4 are shown. Inset (**i**) shows zoomed-in view of bead # 4. **(J)** Average nearest neighbor distances calculated from Neurolucida 360 reconstructions of 90 ± 5 nm axially spaced GATTA bead nano rulers (d = 93.6 ± 2.9 nm, n = 133 beads). Graph represents means with individual GATTA bead measurements shown as dots. **(K)** Average volume of individual GATTA beads calculated from 3D Neurolucida 360 reconstructions of 90 ± 5 nm axially spaced GATTA bead nano rulers (v = 0.0012 ± 0.0001 µm^3^, n = 136 beads). Graph represents means with individual GATTA bead volumes shown as dots.

**Figure S3.**
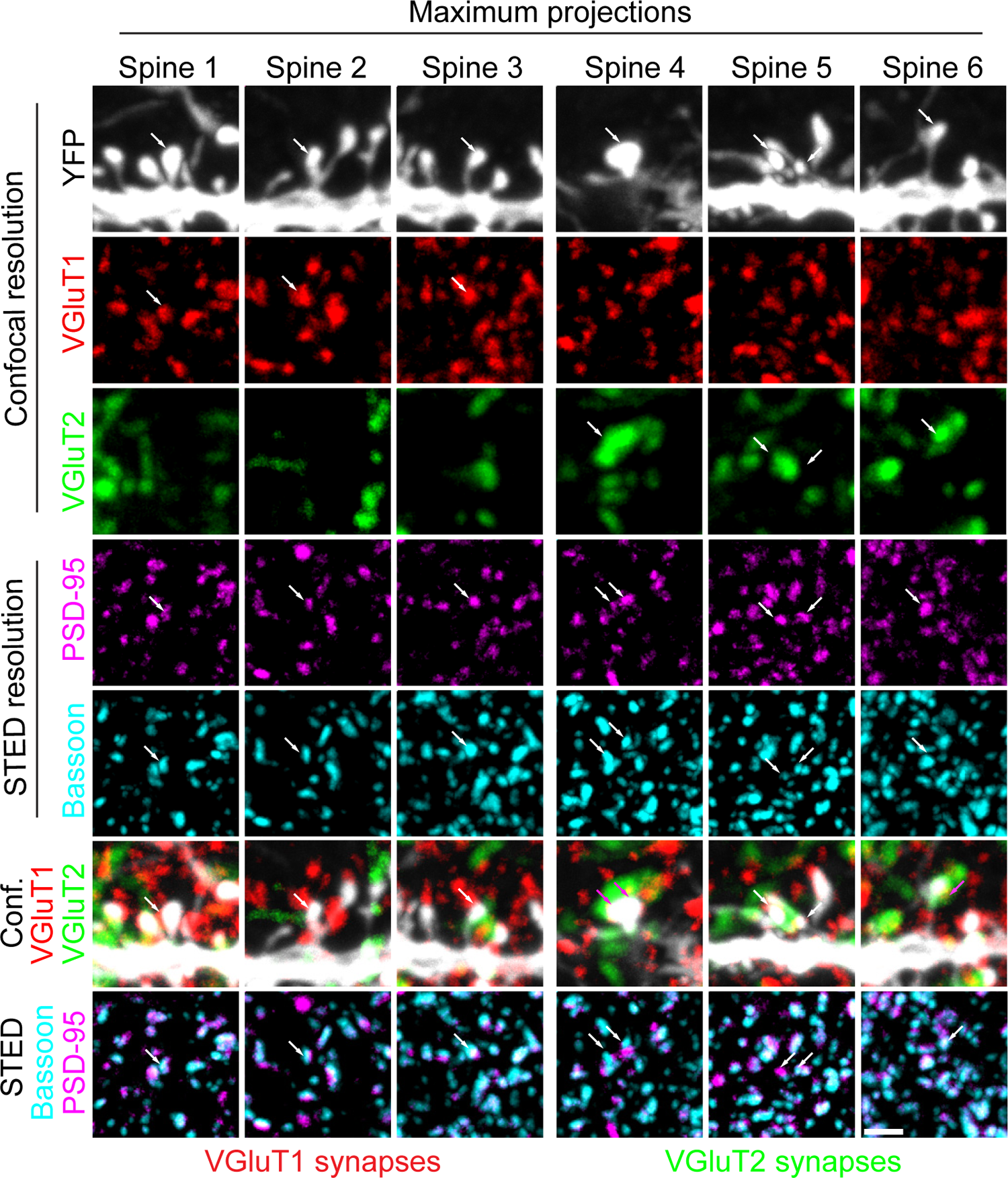
Maximum intensity projection images demonstrate complex distribution of synaptic clusters and spines in stacks through basal dendrites. Representative maximum projection images of 5 channel confocal/tau-STED images of YFP-labeled dendritic spines in basal dendrites of L5 pyramidal neurons from S1 of Thy-1-YFP-H mice. Stacks were the accumulation of 12 individual images in the indicated channels taken at ∼ 160 nm intervals, resulting in a Z-depth of ∼ 2 µm. YFP (white, enhanced by the labeling with GFP antibody), VGluT1 (red, nanobody/Atto 542) and VGluT2 (green, antibody/Alexa Fluor 750) were imaged in confocal resolution. PSD-95 (magenta, nanobody/Abberior STAR 635P) and Bassoon (cyan, antibody/Alexa Fluor 594) were imaged in tau-STED resolution. Arrows indicate single dendritic spines shown in figure 6 that were reconstructed in 3D using Neurolucida 360. The bottom two rows depict merged confocal and tau-STED images demonstrating a large number of clusters found in 2 µm stacks. Scale bar, 1 µm.

**Figure S4.**
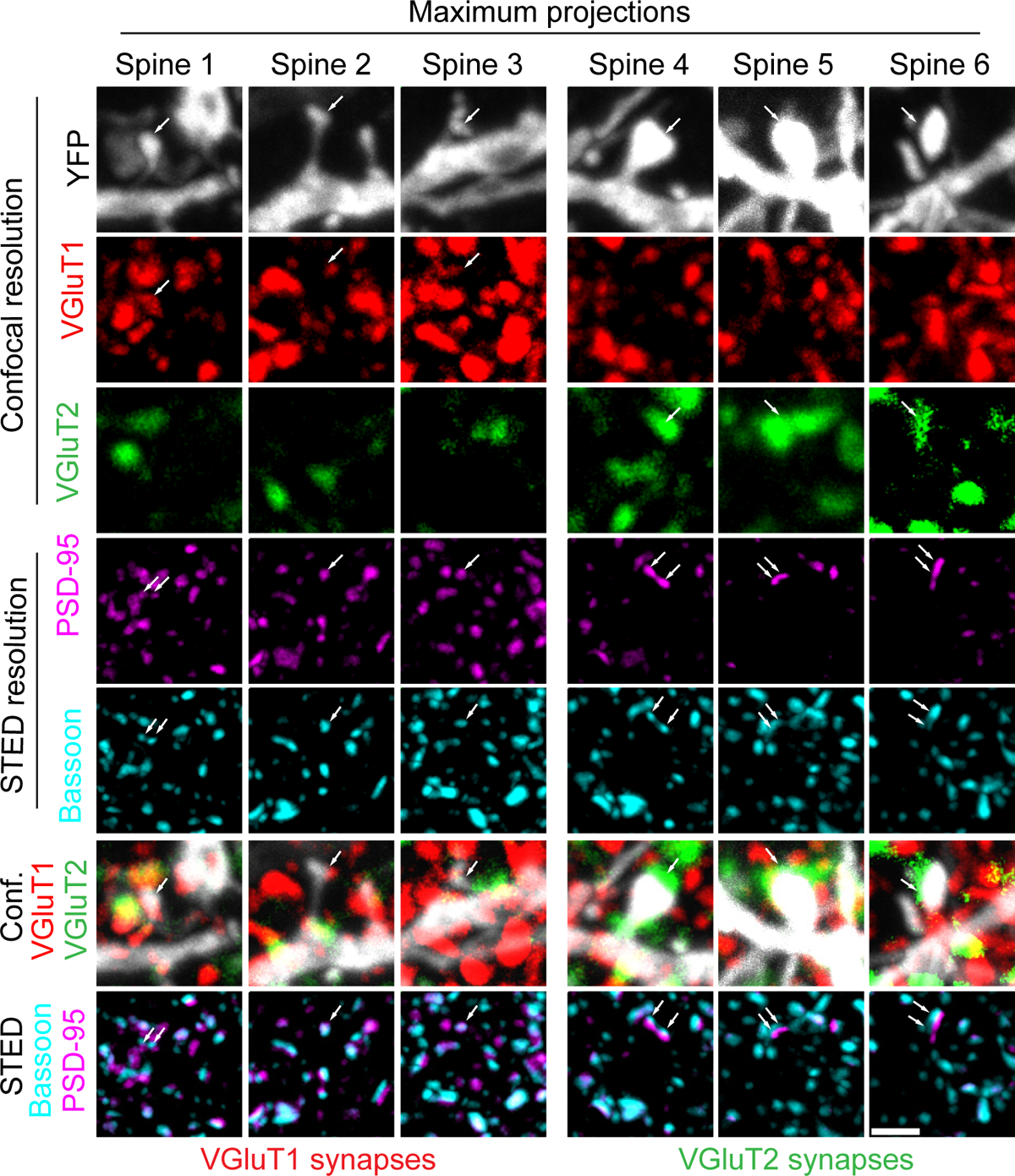
Maximum intensity projection images demonstrate complex distribution of synaptic clusters and spines in stacks through apical dendrites. Representative maximum projection images of 5 channel confocal/tau-STED images of YFP labeled dendritic spines in apical dendrites of L5 pyramidal neurons from S1 of Thy-1-YFP-H mice. Stacks were the accumulation of 17 individual images in the indicated channels taken at ∼ 170 nm intervals, resulting in a Z-depth of ∼ 2.8 µm. YFP (white, enhanced by the labeling with GFP antibody), VGluT1 (red, nanobody/Atto 542) and VGluT2 (green, antibody/Alexa Fluor 750) were imaged in confocal resolution. PSD-95 (magenta, nanobody/Abberior STAR 635P) and Bassoon (cyan, antibody/Alexa Fluor 594) were imaged in STED resolution. Arrows indicate single dendritic spines shown in figure 7 that were reconstructed in 3D using Neurolucida 360. The bottom two rows depict merged confocal and tau-STED images demonstrating a large number of clusters found in 2.8 µm stacks. Scale bar, 1 µm.

## Supplementary video figure legends

**Supplementary Video 1.** 3D reconstruction of confocal-resolved VGluT1 and VGluT2 clusters and corresponding YFP-labeled apical dendrites. Representative 23 x 23 x 2.8 µm reconstructions of confocal channels in apical segments of L5 pyramidal neurons. Movie was generated by 3D-rendering of 17 optical sections spaced at 170 nm from 5-channel confocal/3D-tau-STED imaging of apical dendrites and dendritic spines on L5 pyramidal neurons in S1 of Thy-1-YFP-H mice. YFP-labeled dendrites and spines (white, enhanced by the labeling with GFP antibody), VGluT1 (red, nanobody/Atto 542) and VGluT2 (green, antibody/Alexa Flour 750) were imaged in confocal resolution. Dendrites and clusters were reconstructed using Neurolucida-360 to assign CC and TC synapses to YFP-labeled spines based on their colocalization with VGluT1 or VGluT2, respectively. Still images of this apical dendrite and dendrtic spines are shown in figure 7 and figure S4.

**Supplementary Video 2.** 3D reconstruction of STED-resolved PSD-95 and Bassoon in YFP-labeled apical dendrites of L5 pyramidal neurons. Representative 23 x 23 x 2.8 µm reconstructions of STED channels generated by 3D-rendering of 17 optical sections spaced at 170 nm from 5-channel confocal/3D tau-STED imaging of apical dendrites on L5 pyramidal neurons in S1 of Thy-1-YFP-H mice. Dendrites and dendritic spines labeled with YFP (white, enhanced by the labeling with GFP antibody) were imaged in confocal resolution. PSD-95 (magenta, visualized by nanobody/Abberior Star 635P) and Bassoon (cyan, antibody/Alexa Fluor 594) were imaged in STED resolution. Dendrites and STED-resolved PSD-95 and Bassoon nanoclusters were reconstructed using Neurolucida-360 to determine the nano-organization of aligned pre- and post-synaptic components of active zones and PSDs in VGluT1^+^ or VGluT2^+^ dendritic spines. Still images of this apical dendrite and dendritic spines with associated nanoclusters are shown in figure 7 figure S4.

**Supplementary Video 3.** 3D-rendering of aligned PSD-95 and Bassoon nanoclusters in molecularly defined YFP-labeled dendritic spines. Following the 3D reconstruction of confocal and STED channels, only clusters that colocalized with YFP-labeled dendritic spines were analyzed. Dendritic spines shown in peach color represent synapses receiving VGluT2 input. White spines represent synapses receiving VGluT1 input. Nano-organization of PSD-95 (magenta) and Bassoon (cyan) was determined by rendering tau-STED resolved nanoclusters in VGluT1^+^ and VGluT2^+^ spines. PSD-95 and Bassoon nano-architecture of all peach-colored VGluT2^+^ spines is shown. Volumes for quantification were obtained from 3D-rendered spines and associated PSD-95 nanoclusters. Still images of these dendritic spines are shown in figure 7.

